# Structural and mechanistic insights into Dis3L2–mediated degradation of structured RNA

**DOI:** 10.1101/2025.08.01.668158

**Authors:** Rute G. Matos, Ankur Garg, Susana M. Costa, Patrícia Pereira, Cecília M. Arraiano, Leemor Joshua-Tor, Sandra C. Viegas

## Abstract

The RNase II/RNB family of exoribonucleases is present in all domains of life and includes three main eukaryotic members, the Dis3-like proteins (Dis3, Dis3L1, Dis3L2). At the cellular level, Dis3L2 is distinguished by the unique preference for uridylated RNA substrates and the highest efficiency in degrading double-stranded RNA. Defects in these enzymes have been linked to some types of cancers and overgrowth disorders in humans. In this work, we used the Dis3L2 protein from the model organism *Schizosaccharomyces pombe* (SpDis3L2) to better understand the mechanism of action of Dis3-like exoribonucleases, and to elucidate how single amino acid substitutions in these proteins can affect the biochemical properties of the enzymes, potentially contributing to the molecular basis of the related human diseases.

We determined the crystal structure of SpDis3L2 bound to a U_13_ RNA, in which the protein displays a typical vase-like conformation, accommodating 6 nucleotides of the RNA 3’-end. Furthermore, we constructed two SpDis3L2 protein variants, harbouring single amino acid substitutions mimicking the ones already found in human patients, to test their catalytic activity *in vitro*. We highlight the A756R SpDis3L2 variant, which loses the ability to degrade double-stranded RNA substrates and accumulates intermediary degradation products when degrading single-stranded RNA substrates. As such, A756 seems to be a key residue responsible for the normal cellular function of Dis3L2, specifically regarding its important role in the degradation of structured RNA substrates.

## Introduction

Ribonucleases (RNases) are enzymes that process and degrade all types of RNA, and are critical for the post-transcriptional control of gene expression. The RNase II/RNB family of 3’-5’ processive exoribonucleases cleaves RNA substrates starting from their 3’-end. This family of RNases is characterised by a highly conserved central catalytic RNB domain (named after the *rnb* gene, which encodes the bacterial RNase II) (Frazão et al. 2006), and three other RNA-binding domains – cold-shock domains 1 and 2 (CSD1 and CSD2) and S1. Members of this family can be found in prokaryotes (RNase II and RNase R) and eukaryotes (Dis3– also known as Rrp44, Dis3-like exonuclease 1 (Dis3L or Dis3L1) and Dis3-like exonuclease 2 (Dis3L2)) (Costa et al. 2022).

Dis3, Dis3L1, and Dis3L2 enzymes have already been linked to several human diseases (reviewed in (Saramago et al. 2019; Costa et al. 2022)), such as Multiple Myeloma (Tomecki et al. 2014; Robinson et al. 2018), Cerebellar Medulloblastoma (Palmer et al. 2014), and Perlman Syndrome (Astuti et al. 2012; Higashimoto et al. 2013; Soma et al. 2017; Gowans et al. 2021; Friedman et al. 2023), respectively. Perlman Syndrome is associated with human Dis3L2 gene (*hDIS3L2*) mutations and/or deletions (Wegert et al. 2015; Hol et al. 2022; Friedman et al. 2023). It is a rare overgrowth disease in newborns leading to high infant mortality, which also leads to a higher risk for a kidney cancer known as nephroblastoma or Wilms’ Tumour. hDIS3L2 has also been associated with some types of cancer, like hepatocellular carcinoma (Xing et al. 2019) (LIVER CANCER, ICGC Data Portal, https://dcc.icgc.org/projects/LICA-CN and https://dcc.icgc.org/projects/LINC-JP, accessed April 22, 2024, currently closed) and colorectal cancer (Liu et al. 2017; García-Moreno et al. 2023) (COLORECTAL CANCER, ICGC Data Portal, https://dcc.icgc.org/projects/COCA-CN, accessed April 22, 2024, currently closed). The study of the overgrowth phenotype in Dis3L2-deficient cells in different animal models revealed cellular processes that Dis3L2 might regulate to induce cell proliferation rather than cell differentiation. These include the PI3-Kinase/AKT signalling pathway (Towler et al. 2020), the endoplasmic reticulum (ER)-targeted mRNA translation/ER calcium release (Pirouz et al. 2020), and the expression of the growth-promoting gene *Igf2* (Hunter et al. 2018).

At the cellular level, while Dis3 and Dis3L1 associate with the multiprotein complex RNA exosome to degrade RNA molecules (Tomecki et al. 2010; Fasken et al. 2020), Dis3L2 acts independently in an alternative cytoplasmic pathway linked to 3’-uridylation (Malecki et al. 2013; Chang et al. 2013; Ustianenko et al. 2013; Lubas et al. 2013). Uridylation consists of the addition of untemplated uridines (U) to the RNA 3’-end. This post-transcriptional modification is mediated by terminal uridylyl transferases (TUTases) (reviewed in (Viegas et al. 2015; De Almeida et al. 2018; Yu and Kim 2020)). Dis3L2 preferentially recognises and degrades uridylated RNA molecules, even if they are highly structured. A well-studied example is the processing pathway of uridylated pre-let-7, the precursor of the human let-7 microRNA (miRNA), which is associated with human tumorigenesis (Heo et al. 2012; Zhang et al. 2022).

The exquisite specificity and preference of Dis3L2 for uridylated substrates have been explored in the structure of mouse Dis3L2 (mDis3L2) in complex with a U_14_ RNA that mimics the U-tail of pre-let-7 (Faehnle et al. 2014). The mDis3L2–U_14_ structure revealed a wide network of interactions between specific amino acids of the protein and the uracil bases of the RNA substrate (Faehnle et al. 2014). These interactions were recapitulated with the cryoEM structures of human Dis3L2 in complex with a series of uridylated structured RNA substrates (Meze et al. 2023).

Dis3L2 requires a minimum of 2 single-stranded (ss) nucleotides protruding from the 3’-end of a double-stranded (ds) region to degrade structured RNA substrates (Malecki et al. 2013). However, it has also been shown to degrade dsRNA with blunt ends, albeit to a much smaller extent than with ss overhangs (Lubas et al. 2013). These capabilities make Dis3L2 much more efficient in degrading structured RNA substrates compared to its eukaryotic paralogues Dis3 and Dis3L1. A model for the molecular mechanism of structured RNA degradation by hDIS3L2 was recently proposed based on structures and extensive kinetics analysis of hDIS3L2 in complex with RNA hairpins with progressively shorter 3’-U overhangs (Meze et al. 2023). According to that model, RNA-free hDIS3L2 is in a vase conformation when substrate association occurs, with around 3 binding events before hDIS3L2 starts degrading the ss 3’-overhang of the RNA substrate in a processive manner. The first contacts between the RNA-binding domains of hDIS3L2 and the ds region of the RNA substrate are established when there is a 12-to 11-nt ss-overhang. When the ss overhang reaches 9-to 8-nt, those dsRNA–hDIS3L2 contacts become clashes and force a drastic conformational rearrangement of the CSD domains of the protein. In that new prong conformation, a trihelix linker region in the RNB domain is exposed and works as a wedge to separate the two RNA strands. This way, hDIS3L2 is capable of unwinding the RNA double-strand while a 5-nt ss-portion of the RNA substrate can enter the protein’s narrow tunnel and continue to be degraded (Meze et al. 2023).

For the *Schizosaccharomyces pombe* Dis3L2 (SpDis3L2) protein, some initial biochemical characterisation (Malecki et al. 2013) and an RNA-free structure of Dis3L2 are available (Lv et al. 2015). In this work, we determined the crystal structure of *S. pombe* Dis3L2 (SpDis3L2) in complex with a U_13_ RNA substrate to better understand its basis of RNA recognition, preference for uridylated RNA substrates, and mechanism of RNA degradation.

Furthermore, there is still a need to understand further the Dis3L2 mechanism of action and its association with human diseases. Proteins from this family in eukaryotes are notoriously challenging to work with, especially the human paralogs. Given that *S. pombe* is a well-established model organism and that these proteins are highly conserved, we consider SpDis3L2 a good candidate to perform functional studies. Taking this into consideration, we searched in the literature for mutations in Dis3-like enzymes that have been linked to disease to analyse their impact on Dis3L2 activity. We constructed two SpDis3L2 variants with single amino acid substitutions (C560Y and A756R) and investigated their effect on the catalytic activity using different RNA substrates. Surprisingly, the A756R SpDis3L2 variant showed serious catalytic abnormalities using both ssRNA and dsRNA as substrates. These observations are supported by the crystal structure of SpDis3L2-U_13_ reported here and reflect the importance of this highly conserved residue in Dis3L2 for its mechanism of action and normal cellular function.

## Results & Discussion

### Overall structure of SpDis3L2

To investigate the structural basis of SpDis3L2 specificity for U-tailed RNA substrates, we purified the catalytically dead SpDis3L2(D461N)^ΔN168^ protein and co-crystallised it with a U_13_ oligonucleotide substrate (**Figure S1**). The crystals belonged to space group P4_1_2_1_2 with 2 molecules in the asymmetric unit (ASU) (**Table 1**). Molecule-1 structure lacks N-terminal amino acids 168-173, a loop peptide in CSD1 (amino acids 241-292), and a loop peptide in CSD2 (amino acids 320-328) since the electron density for these regions was not interpretable. However, the rest of the polypeptide chain could be unambiguously traced in the electron density map. Molecule-2 structure lacks N-terminal amino acids 168-180 and a loop peptide in CSD1 (amino acids 225-228 and 241-290), with the rest of the polypeptide chain easily traceable in the electron density map. The two molecules in the ASU are almost identical (**Figure S1C**) (RMSD of 0.72 Å for 570 aligned Cα atoms). Although we used a U_13_ oligonucleotide for crystallisation experiments, only 12 nucleotides (U2-U13) were observed in the electron density map for both molecules. Since the electron density for the CSD1 in molecule-1 was of higher quality compared to molecule-2, we used molecule-1 for our structural analysis.

**Table 1.**
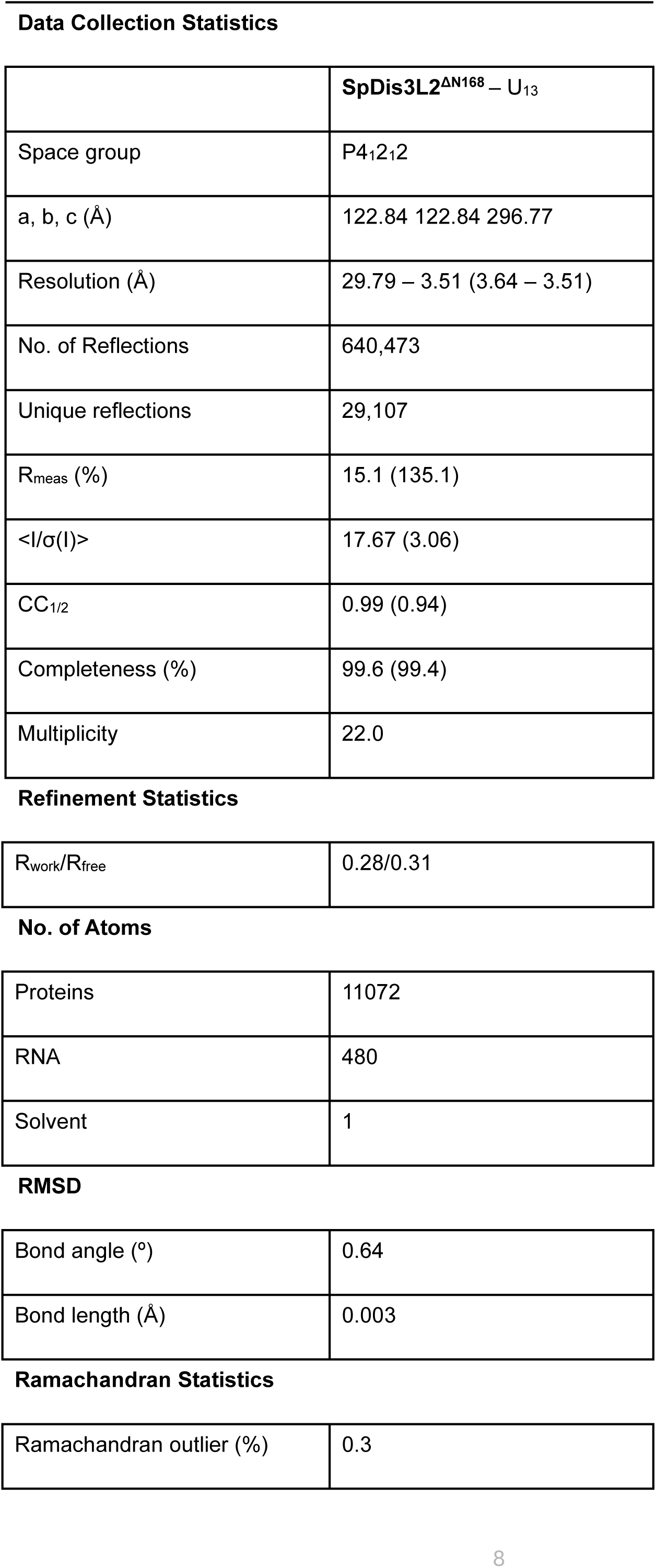
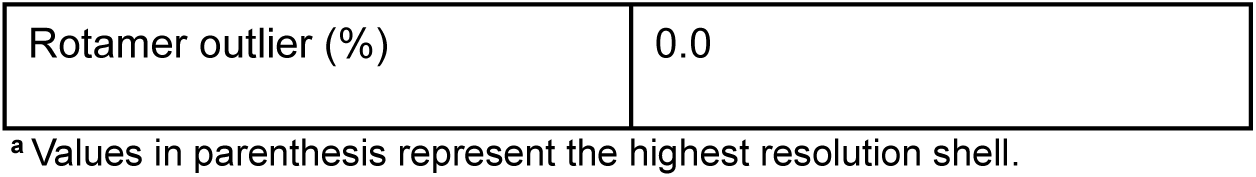
X-ray data collection and refinement statistics for the crystal structure of SpDis3L2 bound to U_13_ RNA.

SpDis3L2 adopts a globular fold with the CSD1, CSD2, and S1 domains assembled on top of the catalytic RNB domain to form a deep funnel-shaped tunnel for RNA binding (**Figure 1A–B**). The CSD1, CSD2, and S1 domains all exhibit positively charged surfaces. CSD1 and S1 interact primarily with the bases and backbone of the 5 nucleotides at the 5’-end (U2-U6), respectively, directing the oligo(U) tail into the tunnel (**Figure 1C**). The catalytic site is buried deep in the RNB domain at the other end of the charged tunnel. The RNB tunnel accommodates 6 nucleotides at the 3’-end (U8-U13), which exhibits a strict RNA geometry (**Figure 1D**), conserved in all Dis3 and Dis3L2 enzymes. Although the U2 nucleotide is positioned next to the positively charged pockets on the CSD2 surface (**Figure S1C**), the very first 5’ U1 nucleotide is not seen in the structure and stays solvent exposed, likely due to lack of interactions with CSD2 or other parts of the enzyme.

**Figure 1.**
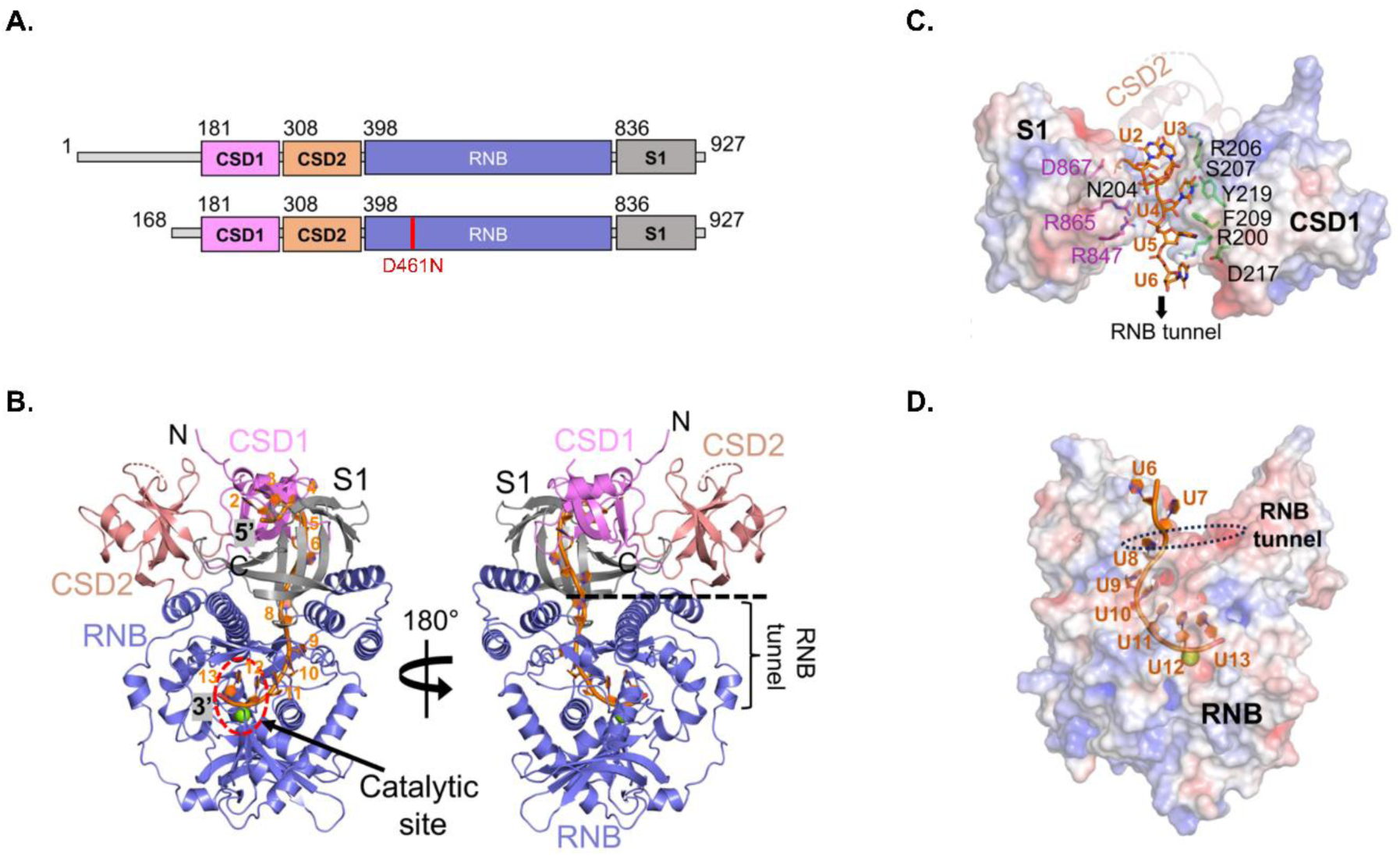
Crystal structure of *S. pombe* Dis3L2. **(A)** Domain architecture of SpDis3L2. The full-length SpDis3L2 (top) and the catalytically inactive SpDis3L2(D461N)^ΔN168^ construct used for crystallization (bottom) are shown as bar diagrams. **(B)** Cartoon representation of SpDis3L2-U_13_ structure in two different views. The CSD1, CSD2, and S1 domains sit atop the RNB domain. The RNA (orange sticks) sits in the RNB tunnel of Dis3L2. The catalytic site is shown with the red dotted circle, marked by the presence of a Mg^2+^ ion (green sphere). The interactions of different U-nucleotides with the CSD1, CSD2, and S1 domains **(C)** and the RNB domain **(D)** are shown. The protein surface is coloured by its electrostatic potential, and RNA is represented as orange sticks. U2 to U5 interact with the CSD1 (amino acids represented in green and designated in black) and S1 (amino acids represented and designated in purple) domains, at the surface of SpDis3L2 **(C)**, while U7 to U13 are buried in the charged RNB tunnel **(D)**.

### Comparison of RNA-free vs. U_13_-bound SpDis3L2

Comparing the SpDis3L2–U_13_ complex structure with the RNA-free SpDis3L2 structure (PDB ID: 4RO1) (Lv et al. 2015) revealed several structural rearrangements upon RNA binding. Although SpDis3L2 has a similar truncation in the RNA-free and the U_13_-bound structures (ΔN170 in RNA-free vs. ΔN168 in U_13_-bound structure), CSD1 was not observed in the RNA-free structure, and it is likely flexible without the RNA. CSD2 was positioned in an open conformation with a widely exposed RNB tunnel in the RNA-free structure (**Figure 2A–B**). In the U_13_-bound structure, CSD2 swings significantly to position itself on top of the RNB domain, adopting a closed conformation (**Figure 2C**) and interacting with the U3, U4, and U5 nucleotides (**Figure S1D**). CSD1, which is flexible in the RNA-free structure, is structured in the U_13_-bound complex and moves with CSD2 to produce the globular SpDis3L2-RNA structure. RNA binding in SpDis3L2 also stabilizes the S1 domain, as we observed a complete S1 domain compared to the RNA-free structure (**Figure 2C**). For the hDIS3L2, on the other hand, ssRNA binding does not change the overall structure much. However, a significant repositioning of the CSD domains does occur in hDIS3L2 upon encountering the double-stranded region of the RNA substrate, albeit in a different manner, which helps it degrade structured RNA substrates, as mentioned above (Meze et al. 2023).

**Figure 2.**
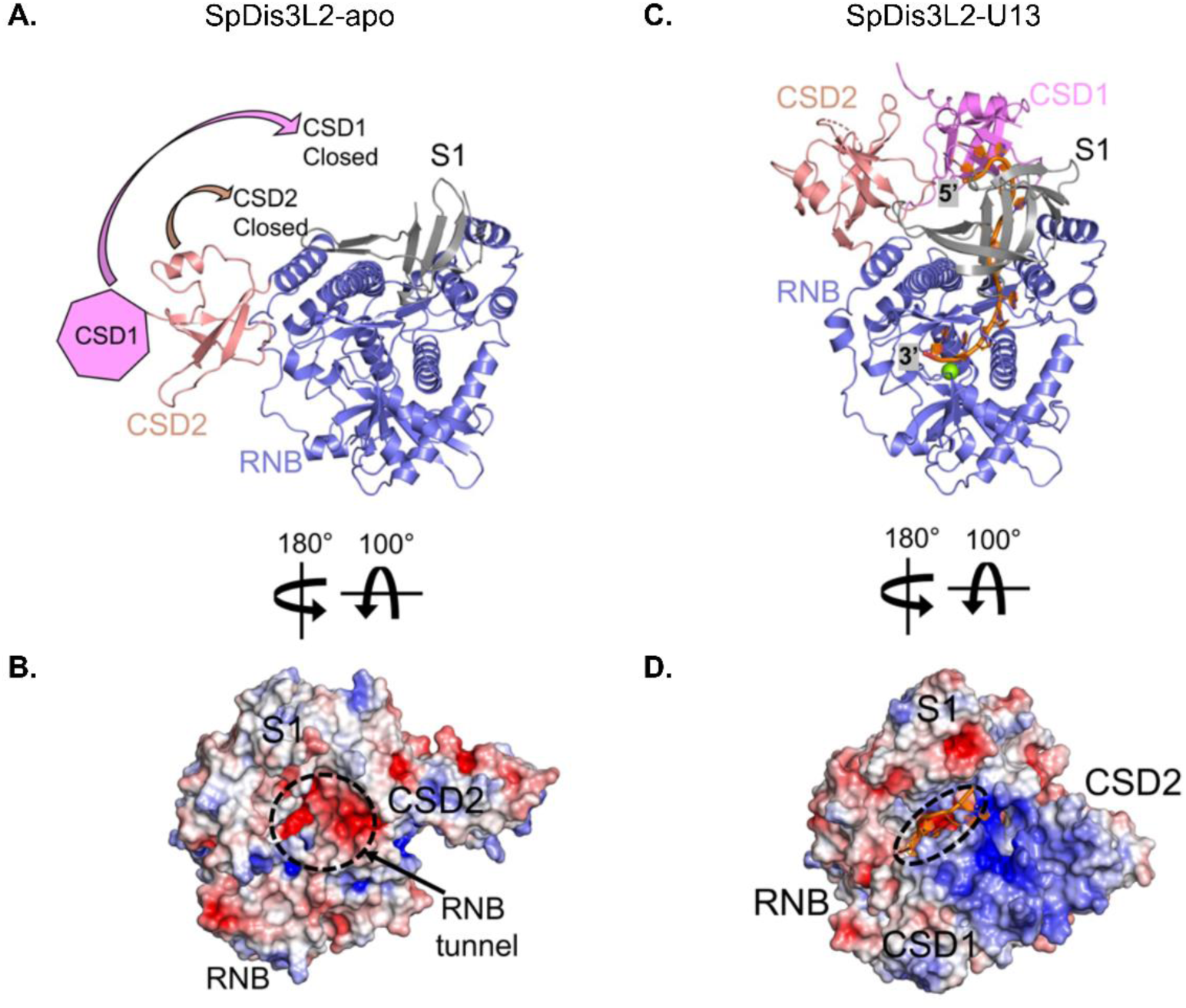
Structural comparison between RNA-free and RNA-bound SpDis3L2. The RNA-free SpDis3L2 structure (Lv et al. 2015) is shown in cartoon representation **(A)** and as a mapping of the electrostatic potential on the protein surface **(B)**, in two different views. CSD1 (pink heptagon) was not observed in the RNA-free structure and its movement along with CSD2 is shown with arrows. Looking at the electrostatic view from the top, the empty charged RNB tunnel is also seen (black dotted circle). The U_13_-bound SpDis3L2 structure is shown in cartoon representation **(C)** and as a mapping of the electrostatic potential on the protein surface **(D)**, in two different views. RNA is represented as orange sticks. The Mg^2+^ ion of the catalytic site is shown as a green sphere. CSD1 and CSD2 significantly reposition on top of the RNB domain upon RNA binding and narrow the tunnel (black dotted ellipse) to encircle RNA. Electrostatic surface potential is displayed in a range from –5 (red) to +5 kT/e (blue) at 0.15M salt concentration.

### Comparison of SpDis3L2–U_13_ vs. mammalian Dis3L2–RNA complexes

Comparing the U_13_-bound SpDis3L2 structure with the mammalian (mouse and human) Dis3L2–RNA complex (Meze et al. 2023; Faehnle et al. 2014) in the vase conformation revealed differences in the relative positioning of the different domains.

The CSD1 domain in SpDis3L2 is positioned on top of the long α15 helix of the RNB domain. CSD1 is rotated and positioned at least ∼10 Å farther away from the α15 helix in RNB as compared to the human DIS3L2, with minor changes in the protein– RNA interactions (**Figure 3A**). There are only small changes in the positioning of CSD2 and the S1 domain relative to the RNB domain between the two enzymes (**Figure 3B–C**).

**Figure 3.**
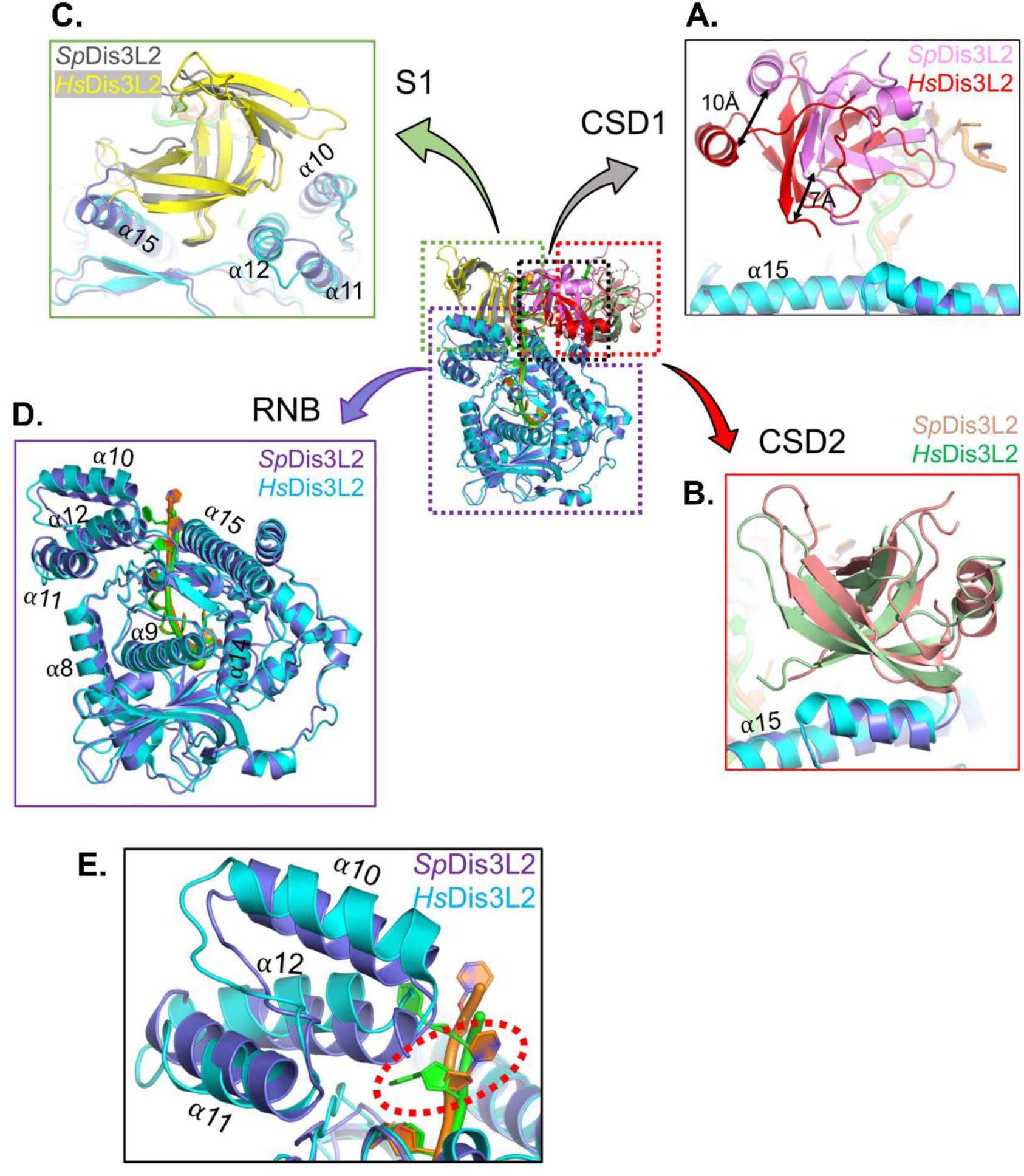
*S. pombe* Dis3L2 (SpDis3L2) vs. human DIS3L2 (HsDis3L2). Structural superposition of RNA-bound SpDis3L2 and human DIS3L2 (HsDIS3L2) CSD1 **(A)**, CSD2 **(B)**, S1 domain **(C)** and RNB domain **(D)** shown as cartoon representations. CSD1 shows significant structural rearrangements (highlighted with black arrows). CSD2 and S1 domains also show subtle structural differences between the two species. The RNB domain fold in the two structures is conserved, but the tri-helical bundle (α10-α12) **(E)** is pushed closer to the rest of the RNB domain, forcing U7 to flip to the other side relative to the subsequent nucleotides (red dotted circle). RNA is represented as orange (in SpDis3L2) and green (in HsDIS3L2) sticks.

Interestingly, there is a subtle change in the RNB domain of SpDis3L2, with its α10-α12 helical bundle (trihelix bundle) pushed slightly downwards in the direction of the active site compared to the human enzyme (**Figure 3D**). In mammalian Dis3L2 (both mouse and human), α12 is positioned on top of the 6^th^ nt from the 3’-end with Ser664 inserted between the 6^th^ and 7^th^ nucleotide from the 3’-end (U7 and U6). However, since α12 is positioned a bit lower in SpDis3L2, it would clash with U7 forcing it to flip to the other side of the backbone and stack on R737 (**Figure 3E**). The trihelix bundle in the RNA-free SpDis3L2 structure is positioned similarly to that structural motif in the RNA-bound structure, indicating that *S. pombe* and mammalian Dis3L2 differ somewhat in the way they position the oligo(U) tail in the RNB tunnel. It should be noted that this nucleotide flips to the other side when comparing the vase and prong conformations of hDIS3L2, though there are also some changes in the backbone as well. These slight structural differences might result in some mechanistic differences in SpDis3L2 catalysis compared to the mammalian enzyme, which should be further investigated.

Notably, the RNA follows a similar path down the catalytic tunnel in all Dis3L2 structures, and the geometry of the 5 nucleotides at the 3’-end in the RNB tunnel is strictly conserved in all Dis3L2 structures.

### Exoribonucleolytic activity of SpDis3L2 variants over ssRNA substrates

In humans, proteins from the RNB-family of enzymes were already shown to be associated with the development of several diseases (Costa et al. 2022; Tomecki et al. 2014; Palmer et al. 2014; Astuti et al. 2012). Mutations in hDIS3L2 were found in patients with Perlman Syndrome, where a specific mutation, C489Y, was found (Astuti et al. 2012). Interestingly, this Cysteine residue is highly conserved in Dis3L2 proteins but not in the other Dis3-like proteins (**Figure S2**). When searching for conserved residues that were only present in Dis3L2 proteins, we noticed that alanine 756 in SpDis3L2 corresponds to a glycine in Dis3 proteins (G766 in hDIS3). Curiously, a mutation in this residue to an arginine was found in a patient with Multiple Myeloma (Tomecki et al. 2014; Robinson et al. 2018). Considering all this, we decided to further study the role of the residues in these two positions in Dis3L2. As such, we constructed two SpDis3L2 variants:

**i)** The C560Y variant mimics the C489Y hDIS3L2 variant found in a human patient with Perlman Syndrome (Astuti et al. 2012). The cysteine (C) in this position is highly conserved within Dis3L2 proteins, but not Dis3 or Dis3L1 proteins (**Figure S2**). The C560 in SpDis3L2–U_13_ structure is positioned towards the base of the RNB domain and points into a hydrophobic pocket formed by I585, H589 and V594 residues (**Figure 4A**). A C560Y mutation would clash with these amino acids and would cause structural repositioning of the C560 surrounding elements in RNB (**Figure 4B**), likely impacting the overall fold and functioning of Dis3L2;
**ii)** The A756R variant mimics the G766R hDIS3 variant found in a human patient with Multiple Myeloma (Tomecki et al. 2014). In this position, an alanine (A) is conserved within Dis3L2-like proteins, while a glycine (G) is found in all the Dis3-like proteins considered in our alignment (**Figure S2**). The conformation of the last 5 nucleotides within the RNB tunnel is highly conserved between Dis3L2 and Dis3 among different species (**Figure 4E**). By examining the SpDis3L2–U_13_ structure, we observe that the conserved A756 is located on a structured loop in the RNB domain, positioned under the trihelix bundle and pointing towards the ribose C4’ atom of the 5^th^ nucleotide from the 3’-end (U8) in the RNB tunnel, forming van der Waal interactions with it (**Figure 4C**). A similar interaction is also observed in human DIS3L2. Upon the A756R substitution, since arginine (R) is larger than alanine (A), it will likely clash with that nucleotide and might disrupt RNA translocation in the tunnel during catalysis (**Figure 4D**);
**iii)** As a control, we used the D461N variant, which is known to be catalytically inactive (Malecki et al. 2013).

**Figure 4.**
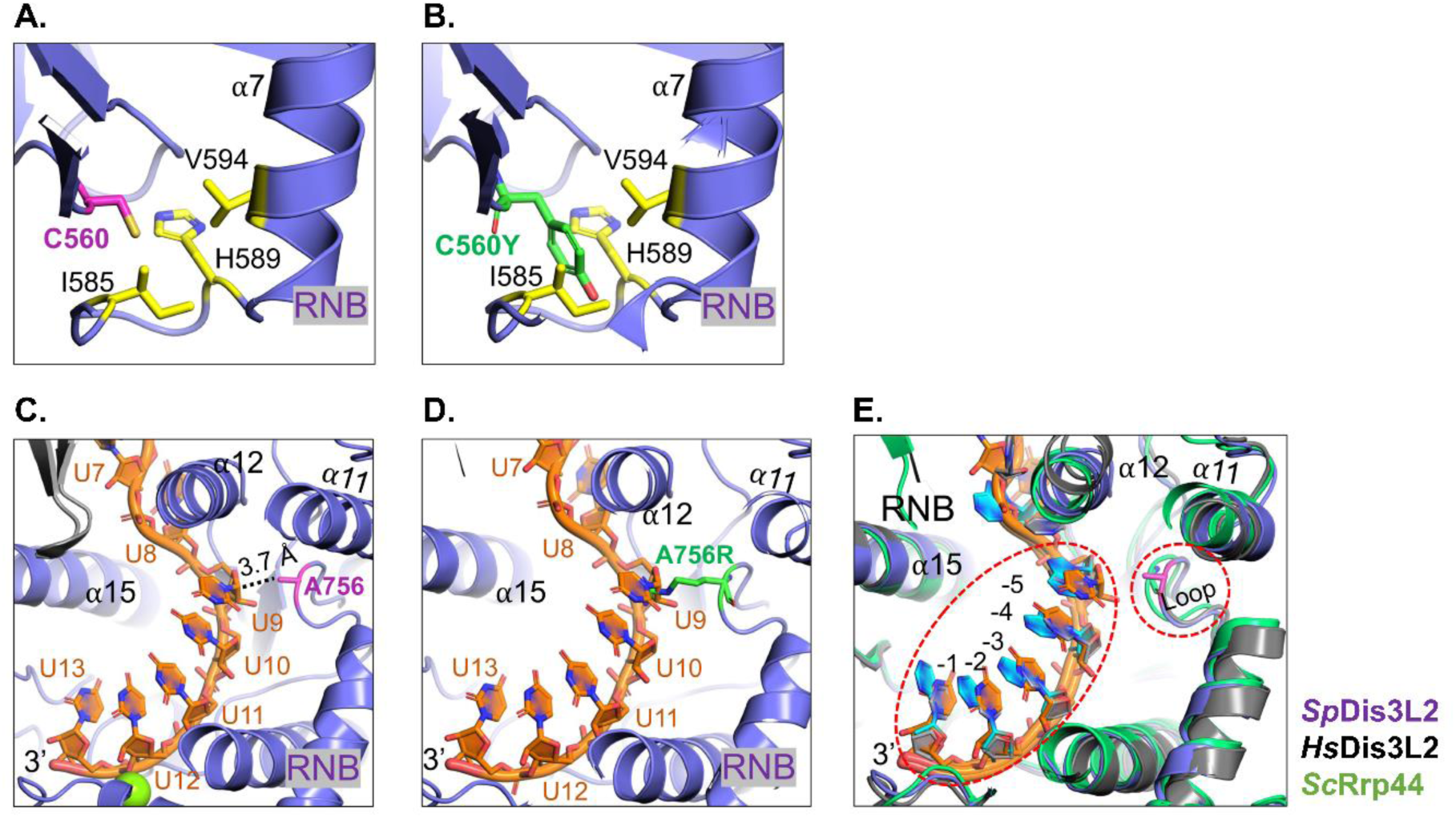
Close-up view of the position of the C560 and A756 residues in the SpDis3L2-U_13_ structure. The RNB domain is shown in blue. α10-α12 constitute the trihelix bundle motif. The Mg^2+^ ion of the catalytic site is shown as a green sphere. RNA is represented as orange sticks. A close-up view into the **(A)** C560 (in magenta) and **(B)** modelled C560Y (in green) interaction with the surrounding hydrophobic residues in the RNB. **(C)** A close-up view of the RNB tunnel highlighting the van der Waals interaction between A756 and the U9 nucleotide. A756 is represented as a magenta stick. The α12 from the trihelix bundle seems to support RNA bending in the RNB tunnel. Different nucleotides down to the 3’-end of the oligo(U) substrate are shown, which adopt a geometry highly conserved in other Dis3 and Dis3L2 proteins. **(D)** A close-up view of the RNB domain with the A756R mutation modelled in the structure. The arginine (R) at this position may establish an additional interaction with U9, thereby impacting RNA substrate processing. **(E)** Superposition of different Dis3L2 and Dis3 structures showing a highly conserved RNA geometry in the RNB tunnel. The positioning of the loop containing A756 (red dotted circle on the right) is also conserved among different structures.

WT, D461N, C560Y, and A756R SpDis3L2 recombinant proteins were overexpressed in *E. coli* and successfully purified with a comparable degree of purity.

To investigate whether the single amino acid substitutions in SpDis3L2 affect its catalytic activity, we conducted exoribonucleolytic activity assays. We used either single-or double-stranded (ss or ds, respectively) synthetic RNA substrates (**Table S3**) to mimic both types of physiological targets that the enzyme is able to degrade (Ustianenko et al. 2016; da Costa et al. 2019; Zhang et al. 2022).

To increase the stability of the protein, in the present study, we constructed a new vector to express SpDis3L2 with a different tag (His-SUMO tag instead of GST tag as in (Malecki et al. 2013), **Table S2**). As such, we re-validated the WT SpDis3L2 protein’s activity. To confirm the size of the end-products, we used an ss poly(U) substrate and either the purified WT SpDis3L2 or RNase II from *E. coli* (EcRNase II). EcRNase II was purified as described in (Frazão et al. 2006). Degradation products of EcRNase II worked as a size marker since it is well-established that this protein produces 4-nt end-products (Frazão et al. 2006; Amblar et al. 2006). We confirmed that the purification of the new WT SpDis3L2 rendered a protein that produces 3-nt end-products (**Figure S3**), as reported previously (Malecki et al. 2013).

To confirm the preference of WT SpDis3L2 for uridylated substrates, we tested its exoribonucleolytic activity on three ssRNA substrates simultaneously: Adh1 (no U-tail at its 3’-end), Adh1-U_4_ (3’-4U-tail), and Adh1-U_16_ (3’-16U-tail). The *Adh1* transcript is known to be uridylated in *S. pombe* (Rissland and Norbury 2009) and was proven to be a substrate of SpDis3L2 (Malecki et al. 2013). We were able to confirm that WT SpDis3L2 prefers longer U-tails as RNA substrates, as Adh1-U_16_ appears to be degraded faster than Adh1-U_4_, and Adh1-U_4_ is degraded faster than Adh1 (**Figure S4A**). We further quantified the disappearance of each full-length substrate along the reaction time to confirm the observed preference trend (**Figure S4B**). In a physiological context, TUTases usually add mono/di(U) tails during the normal biogenesis and/or maturation of certain classes of RNAs. On the other hand, longer oligo(U) tails are added to transcripts to mark them for degradation by Dis3L2 (Viegas et al. 2015) (e.g., up to 7U found in *adh1* transcripts in *S. pombe* (Malecki et al. 2013), up to 15U found in *GADD45A* transcripts with premature translation-termination codons (PTCs) (da Costa et al. 2019), 10-30U found in pre-let-7 (Heo et al. 2012)). Our observations regarding the preference of SpDis3L2 for longer U-tails are consistent with these findings.

To assess the role of the specific amino acid residues of SpDis3L2 in the preference for uridylated RNA substrates, we tested the exoribonucleolytic activity of the SpDis3L2 variants in the presence of two ssRNA substrates simultaneously: Adh1 (non-uridylated) and Adh1-U_4_ (uridylated). As a negative control, we also tested the catalytically inactive D461N SpDis3L2 protein that, as expected, did not show any catalytic activity (**Figure 5**) (Malecki et al. 2013). Both C560Y and A756R SpDis3L2 variants were active, since it was possible to observe the substrates disappearing over time, with increasing accumulation of end-products (**Figure 5**). They also degraded Adh1-U_4_ faster than Adh1, thus showing a preference for uridylated RNA substrates. However, this preference seemed less marked than that of the WT SpDis3L2 (**Figure 5**). The C560Y SpDis3L2 variant presented a similar degradation pattern to the WT enzyme, while the A756R SpDis3L2 variant revealed an accumulation of intermediate degradation products (**Figure 5**). Based on the SpDis3L2–U_13_ structure (**Figure 4B**), the A756R substitution might disrupt normal RNA translocation, causing RNA to frequently dislodge from the enzyme during ssRNA degradation, and, consequently, lead to the release and accumulation of RNA species of varying lengths over time.

**Figure 5.**
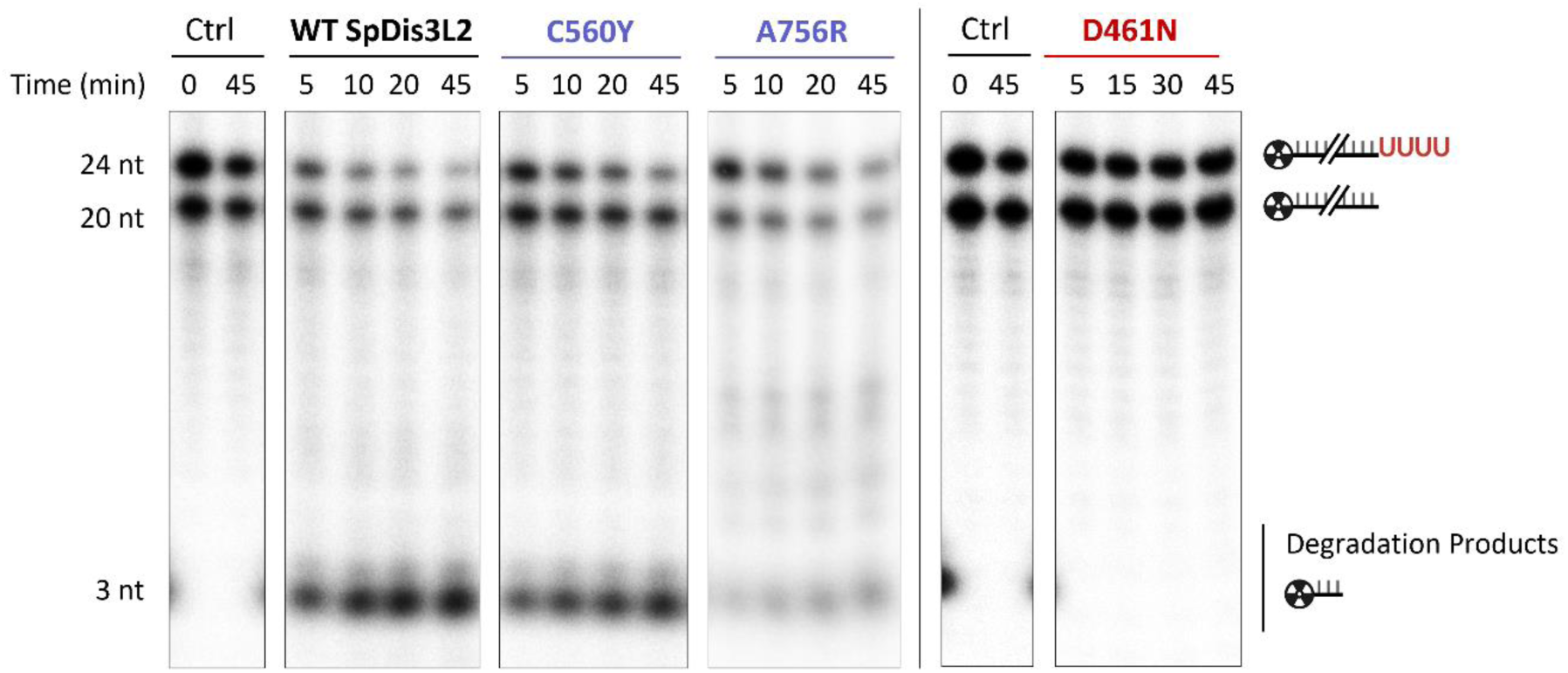
Representative assays of exoribonucleolytic activity of SpDis3L2 wild-type and variants on two different ssRNAs, illustrating their substrate preferences. Adh1 (20 nt) and Adh1-U_4_ (24 nt) RNA substrates were simultaneously incubated for 45 minutes at 30°C with each SpDis3L2 variant (indicated on the top) and the wild-type (WT) protein at the same concentration. Control reactions (Ctrl) were incubated for the same time without any enzyme. Reactions were stopped at the time points indicated above each lane and analysed on a 7 M urea/20% PAA denaturing gel. The sizes of the molecules are depicted alongside the gel in line with the corresponding bands.

### Exoribonucleolytic activity of SpDis3L2 variants over a dsRNA substrate

In eukaryotic cells, Dis3L2 targets a wide range of RNA classes, and it has been implicated in the decay of highly structured non-coding RNAs (ncRNAs) (Ustianenko et al. 2016; Reimão-Pinto et al. 2015). Aberrant ncRNAs usually have a stable double-stranded region near the oligo(U) tail (Ustianenko et al. 2016). Moreover, in comparison with the other eukaryotic ribonucleases of the RNB family (Dis3 and Dis3L1), Dis3L2 is the most efficient in the degradation of structured RNAs *in vitro* (Lubas et al. 2013; Meze et al. 2023).

To evaluate whether SpDis3L2 variants maintained the ability to degrade structured RNA substrates, we incubated them with dsAdh1-U_16_. This substrate results from the hybridisation of ssAdh1-U_16_ with the complementary sequence of Adh1 (asAdh1). This duplex has a 20-nt double-stranded region followed by a 16U 3’-ss-overhang. We confirmed the complete hybridisation of Adh1-U_16_ and asAdh1 to form dsAdh1-U_16_ in a 20% PAA non-denaturing gel (**Figure 6A**).

**Figure 6.**
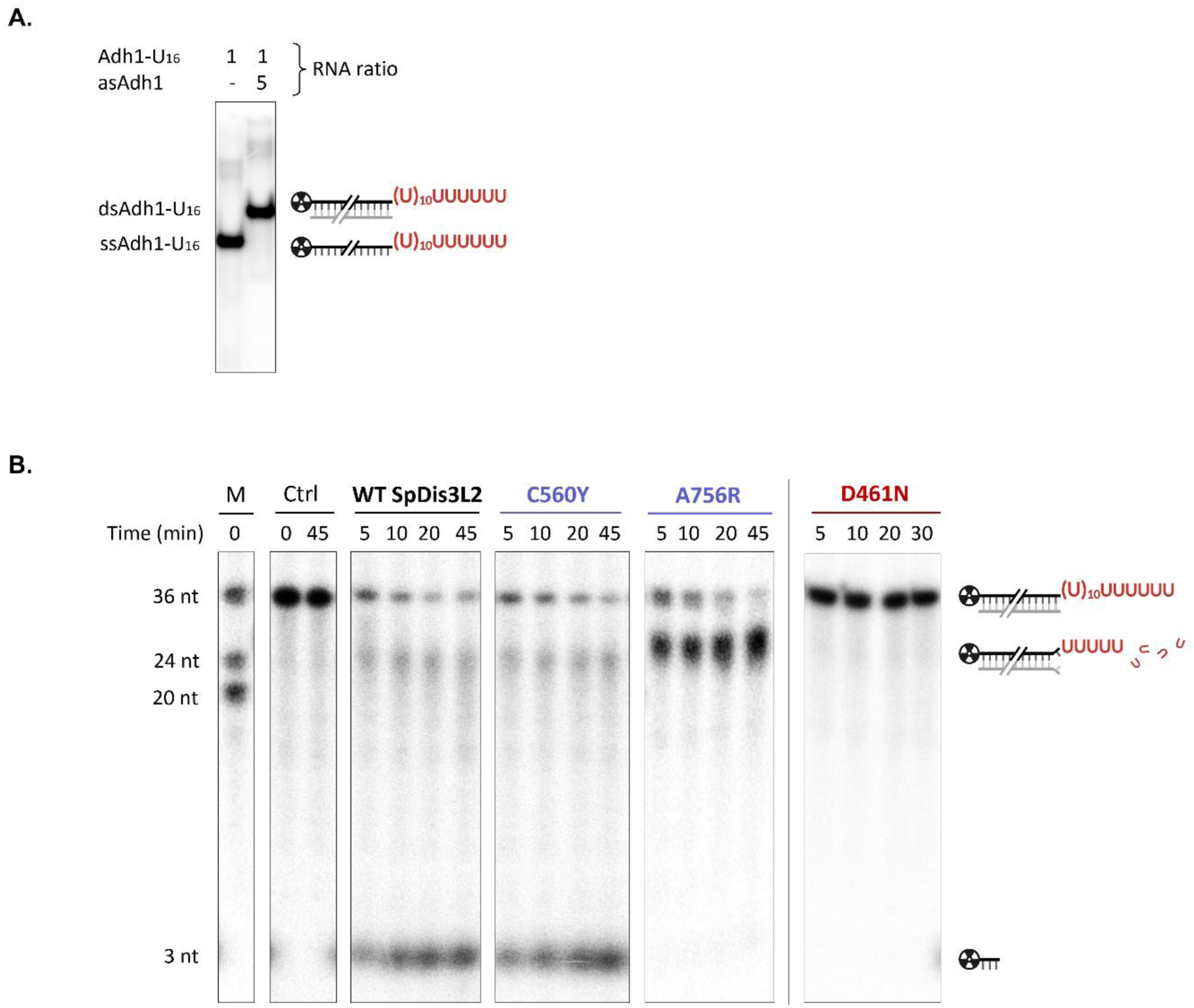
(A) Formation of the dsAdh1-U_16_ substrate. 20% PAA non-denaturing gel of Adh1-U_16_ (ssRNA) and dsAdh1-U_16_ (dsRNA). dsAdh1-U_16_ results from the hybridization of Adh1-U_16_ (labelled; represented in black and U-tail in red) and asAdh1 (the complementary molecule to Adh1, unlabelled; represented in grey) in a 1:5 molar ratio. **(B) Representative assays of exoribonucleolytic activity of SpDis3L2 wild-type and variants on dsAdh1-U**_16_ **RNA substrate.** Labelled Adh1-U_16_ was hybridised with the complementary unlabelled asAdh1 to form dsAdh1-U_16_. The dsAdh1-U_16_ RNA substrate was incubated at 30°C with each SpDis3L2 variant (indicated on the top) and the wild-type (WT) protein at the same concentration. Control reactions (Ctrl) were incubated for the same time without any enzyme. Reactions were stopped at the time points indicated above each lane and analysed on a 7 M urea/20% PAA denaturing gel. The sizes of the molecules are depicted alongside the gel in line with the corresponding bands. ‘M’ indicates the size marker lane with a sample of the three ssRNAs: Adh1 (20 nt), Adh1-U_4_ (24 nt) and Adh1-U_16_ (36 nt).

C560Y SpDis3L2 variant was able to degrade the structured substrate up to a 3-nt end-product and presented a similar degradation pattern to the WT enzyme (**Figure 6B**). We observed an accumulation of large intermediate degradation products with a molecular weight of around ∼ 24-25 nt, which corresponds up to a ∼ 4-5 nt ss overhang before the beginning of the double-stranded region of dsAdh1-U_16_. This might mean that SpDis3L2 (WT and the C560Y variant) slow down as the double-stranded region of dsAdh1-U_16_ approaches the entrance of the RNB tunnel to the inside of the enzyme. As a result, a small fraction of the RNA molecules is being released. Nonetheless, the WT and C560Y SpDis3L2 proteins are able to overcome the beginning of the double-stranded region and continue degrading the RNA molecule until the final end-product of 3 nt. This result is in accordance with the model recently proposed for the degradation of structured RNA by hDIS3L2 (Meze et al. 2023). Accordingly, after a drastic conformational change, when the RNA has a 4-5 nt ss overhang, the RNA duplex engages with the trihelix bundle in the RNB domain near the entrance of the RNB tunnel. At that point, hDIS3L2 starts the unwinding of the double-stranded region and simultaneous degradation of the ss-portion entering the RNB narrow tunnel. We speculate that, for SpDis3L2, a similar structural rearrangement occurs. This would expose the RNB trihelix bundle thus allowing the unwinding of the ds-portion of the RNA substrate.

Regarding the A756R SpDis3L2 variant (**Figure 6B**), it is able to degrade the 3’-ss-overhang of the dsAdh1-U_16_ RNA substrate, but it stalls a few nucleotides before the double-stranded region, releasing products of around ∼ 24-25 nt. This suggests that the A756R protein is unable to unwind the double-stranded region of the dsAdh1-U_16_ substrate, thus stalling there without proceeding with the degradation.

Considering the model for the degradation of structured RNA by hDIS3L2, and that the A756 residue in SpDis3L2 is in the vicinity of the RNB trihelix bundle (**Figure 4A**), we can speculate that the A756R substitution in SpDis3L2 might perturb the normal positioning of the trihelix bundle or even prevent the correct structural rearrangements necessary for dsRNA degradation to occur. Therefore, it would impair dsRNA unwinding and prevent the reaction from proceeding. This is consistent with our experimental data showing that the A756R SpDis3L2 variant stalls when it encounters dsRNA, contrary to what is seen for the WT SpDis3L2 protein, that can continue the degradation of the dsAdh1-U_16_ substrate until the end (**Figure 6B**).

### The implications of the A756R substitution in SpDis3L2

The results obtained with the A756R SpDis3L2 variant, namely the accumulation of intermediate degradation products when digesting ssRNA substrates (**Figure 5**), and the inability to degrade dsRNA substrates (**Figure 6**), led us to test whether there was a change in the affinity for RNA substrates.

For that, we performed an EMSA of WT vs. A756R SpDis3L2 proteins with the ss Adh1-U_16_ substrate. Interestingly, the A756R SpDisL2 variant seemed to have a higher affinity for the Adh1-U_16_ RNA when compared to the WT protein, since a lower concentration of the A756R protein was needed to bind the same amount of RNA (**Figure 7**). As arginine (R) has a positively charged side chain, in contrast to alanine (A), additional ionic interactions could be formed with the RNA backbone, thus increasing the affinity of the A756R SpDis3L2 variant to RNA.

**Figure 7.**
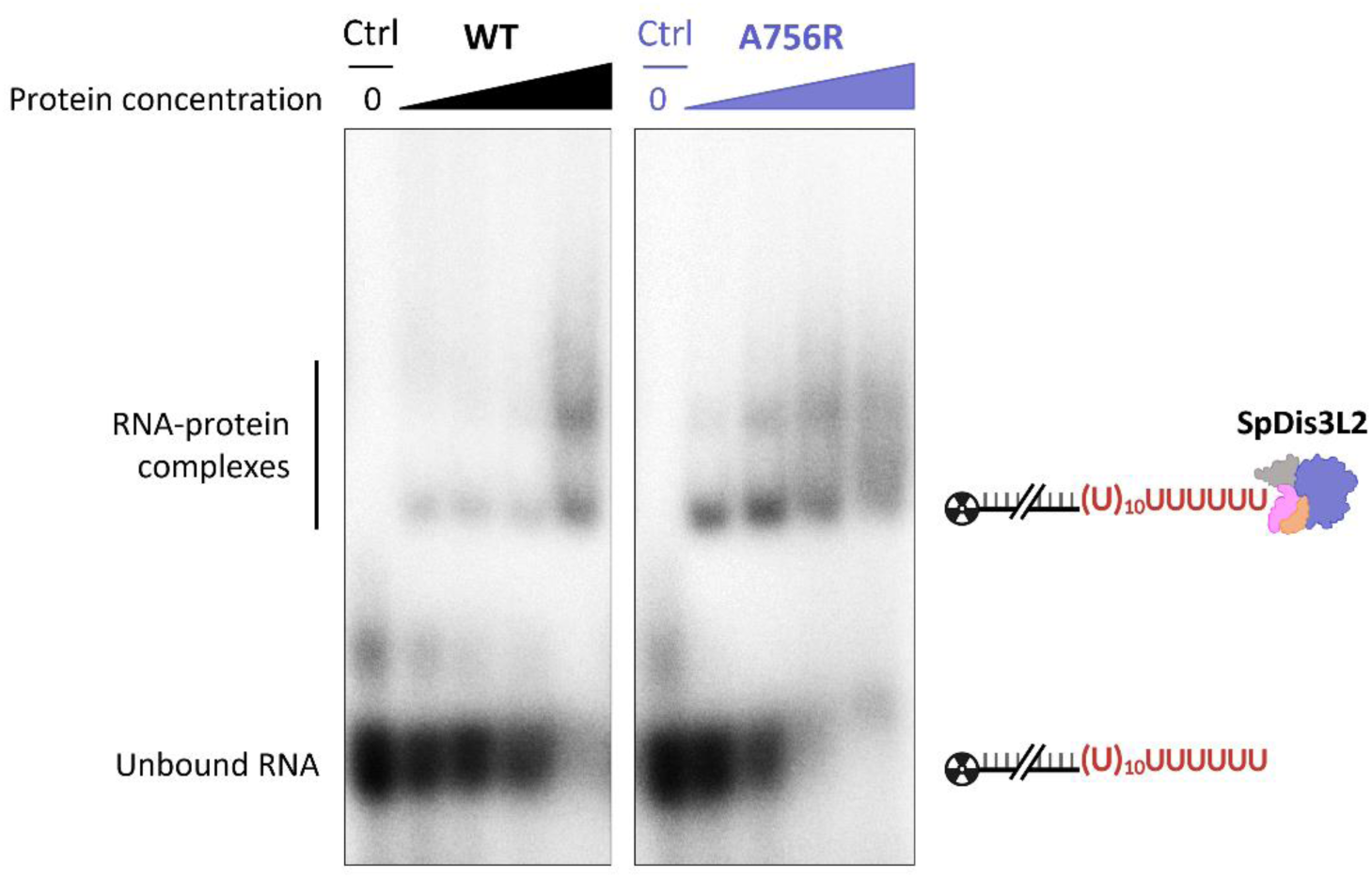
RNA-binding ability of SpDis3L2 wild-type and A756R variant. 10 nM of Adh1-U_16_ ssRNA substrate were incubated for 10 minutes at 30°C with increasing concentrations of each SpDis3L2 variant (indicated on the top): 0 nM (Control, Ctrl), 50 nM, 100 nM, 250 nM, 500 nM. Reaction products were resolved in a 5% PAA non-denaturing gel. Similar results were obtained with proteins from two independent purifications.

It is worth noting that, during the degradation of an RNA substrate by an RNase, there is a fine balance between RNA catalysis and the translocation of the RNA substrate towards the catalytic center. So, we can speculate that the tighter RNA binding of the SpDis3L2 A756R variant might disturb this balance. Furthermore, if a continuous movement of the RNA going into the RNB tunnel would be required for an efficient function of the trihelix bundle as a wedge to separate the two strands of the dsRNA, it would be much more difficult for A756R SpDis3L2 variant to proceed with the degradation of dsRNA substrates. This may provide additional insight into why the A756R variant stalls when encountering double-stranded regions in the RNA substrates (**Figure 6**).

## General conclusions & Final remarks

In this work, we determined the crystal structure of SpDis3L2 bound to an RNA substrate. It revealed that a conformational change of the CSD domains occurs upon binding of the RNA. The trihelix bundle motif in the RNB domain is positioned similarly both in RNA-free and RNA-bound SpDis3L2 structures, but it is pushed downwards in comparison with the mammalian structures (mDis3L2 and hDIS3L2). These peculiarities might result in some differences in the SpDis3L2 catalytic mechanism, which should be further investigated.

We also designed, cloned, and purified C560Y and A756R SpDis3L2 variants to study the impact of single amino acid substitutions on the catalytic performance of the enzyme. Those modifications were selected for their importance in human diseases. The C560Y SpDis3L2 variant showed a slightly impaired activity when degrading ssRNA and dsRNA substrates. However, the A756R SpDis3L2 variant substantially differed from the WT SpDis3L2. Notably, for the degradation of ssRNA substrates (Adh1 and Adh1-U_4_), this variant presented a less marked preference for uridylated RNA and accumulated intermediate products. We postulate that the A756R substitution leads to a disruption of RNA translocation in the RNB tunnel during catalysis, thus causing the release of larger RNA species. Concerning the degradation of dsRNA substrates (dsAdh1-U_16_), the catalytic activity of the A756R variant was almost completely abolished when the double-stranded portion of the substrate reached the enzyme. We showed that the A756R variant has a higher affinity for the Adh1-U_16_ RNA than the WT protein. Indeed, we hypothesize that the A756R substitution is responsible for that tighter binding to the RNA substrate and, it could also possibly affect structural rearrangements of the enzyme during the degradation of dsRNA substrates, or the proper positioning/functioning of the trihelix bundle as a wedge to unwind secondary structures, causing the enzyme to stall when it approaches double-stranded regions of the RNA substrates.

The A756R SpDis3L2 variant was designed to mimic the G766R mutation occurring in the corresponding position of hDIS3, which is associated with Multiple Myeloma (Tomecki et al. 2014). A previous study also reported that the hDIS3^G766R^ mutation compromises the efficiency of the DIS3 enzyme, generating large intermediate products for ssRNA substrates, and stalling when encountering a dsRNA substrate (Tomecki et al. 2014). We observed similar defects in RNA processing for the SpDis3L2^A756R^ variant in our study. These observations indicate that a single substitution in this highly conserved residue causes significant defects in the activity of these enzymes. Here, we present *in vitro* evidence, but direct *in vivo* experiments should also be conducted in the future to further assess the impact of A756R SpDis3L2 variant in cell functioning.

Dis3 operates in the context of the RNA exosome, where the combined machinery allows for unwinding structured RNA (Tomecki et al. 2010; Fasken et al. 2020). However, Dis3L2 acts independently in an alternative cytoplasmic pathway linked to 3’-uridylation, namely over highly structured substrates (Malecki et al. 2013; Lubas et al. 2013). The enzyme was seen to be much more effective in degrading dsRNA substrates than the Dis3 or Dis3L1 homologues *in vitro* (Lubas et al. 2013; Meze et al. 2023) and this property should be related to Dis3L2 cellular function and substrates. Therefore, in this work we present biochemical and structural data that contribute to the identification of a residue that seems to be responsible for the degradative activity of Dis3L2 over structured substrates, which is of utmost importance to understand its mechanism of action.

## Materials & Methods

### Protein crystallisation and structural determination

The gene encoding the catalytically dead mutant D461N of SpDis3L2 (Malecki et al. 2013) was subcloned into a pQlinkH (Scheich et al. 2007) vector via Sequence and ligation-independent cloning (SLIC), truncated at the N-terminus [ΔN168] (SpDis3L2^ΔN168^), with a TEV cleavable N-terminal 7xHis tag, and extra C-terminal Asn-Ala-Ala-Ala amino acids (primers listed in **Table S1**). All constructs were verified by DNA sequencing (Genewiz, USA).

The plasmid harbouring SpDis3L2^ΔN168^(D461N) (**Table S2**) was transformed into *Escherichia coli* BL21 (DE3) RIPL cells (Agilent) to produce the recombinant protein. *E. coli* cultures were grown in Terrific Broth (TB) medium (Thermo Fisher Scientific) supplemented with 100 μg/ml ampicillin at 37°C. When cultures reached an OD_600_ of ∼1.2, protein expression was induced by adding 0.5 mM IPTG. Cells were incubated at 18°C for 16 h before harvesting by centrifugation.

Cells were harvested in lysis buffer (25 mM Tris pH 8.0, 500 mM NaCl, 5 mM ß-Me, 10 mM Imidazole, 5 U/ml Benzonase, and 1 μg/ml Lysozyme), supplemented with PI cocktail (Pepstatin, Leupeptin, PMSF, Benzamidine, and Aprotinin), and lysed by sonication. Lysates were clarified by ultracentrifugation at 40,000 rpm for 1 h.

Using the cleared lysate, Ni-NTA affinity chromatography was performed. 7xHis-Dis3L2^ΔN168^ protein was eluted in Ni-NTA elution buffer (25 mM Tris pH 8.0, 100 mM NaCl, 5 mM ß-Me, and 250 mM Imidazole). The His tag was removed by TEV protease cleavage overnight at 4°C. The cleaved Dis3L2^ΔN168^ protein was diluted to reduce the NaCl concentration to 50 mM using dilution buffer (25 mM Tris pH 8.0, and 5 mM ß-Me), and it was loaded onto a HiTrap Heparin-HP column (Cytiva) pre-equilibrated with ion exchange buffer (25 mM Tris pH 8.0, 50 mM NaCl, and 5 mM ß-Me). Dis3L2^ΔN168^ protein was eluted using a NaCl linear gradient from 50 to 1000 mM. The peak fractions were further subjected to one round of anion exchange using HiTrap Q-HP column (Cytiva), and Dis3L2^ΔN168^ protein was similarly eluted using a NaCl linear gradient. The fractions containing Dis3L2^ΔN168^ were pooled and concentrated. Concentrated protein samples were injected into a Superdex 200 increase 16/60 column (Cytiva) pre-equilibrated with SEC buffer (20 mM Tris pH 8.0, 100 mM NaCl, 1 mM DTT, and 1 mM MgCl_2_). Purified Dis3L2^ΔN168^(D461N) protein was concentrated to 10 mg/ml before flash-freezing in liquid N_2_.

Purified Dis3L2^ΔN168^(D461N) protein was incubated with a U_13_ RNA substrate (**Table S3**) in a 1:1.2 molar ratio at room temperature (RT) for 30 min. Crystals of the Dis3L2^ΔN168^–U_13_ complex were obtained using the hanging drop vapour-diffusion method by mixing 8.5 mg/ml protein–RNA complex with reservoir buffer (12% PEG 4000, 100 mM HEPES pH 7.5, and 100 mM sodium acetate) to a final volume of 0.3 μl. The crystals were harvested and cryo-protected using reservoir buffer supplemented with 20% ethylene glycol, and flash-frozen in liquid N_2_.

X-ray diffraction data were collected at beamline 17-ID-2 (λ = 0.9793 Å) at NSLS-II at Brookhaven National Laboratory. Data were processed using XDS (Kabsch 2010), and the phase problem was solved by molecular replacement (MR) using the AlphaFold v.2.0 (Jumper et al. 2021) predicted model as a template in Phaser (McCoy et al. 2007). Model building was done using Coot (Emsley et al. 2010), and refinement was performed in Phenix (Adams et al. 2010). The atom contacts and model geometry were validated using the MolProbity server (Chen et al. 2010), and structure figures were generated using PyMOL v.2.5.5 (Schrodinger, LLC). The APBS tool (Baker et al. 2001) was used to calculate the electrostatic surface potential. The data collection and refinement statistics for the structure are summarized in **Table 1**.

### Protein sequence alignment

A multiple sequence alignment was constructed using T-Coffee v.11 (Notredame et al. 2000; Di Tommaso et al. 2011), available at http://tcoffee.crg.cat/apps/tcoffee/do:regular, and the amino acid sequences of the homologues of the RNase II/RNB family present in the following eukaryotic species: *Schizosaccharomyces pombe* – SpDis3 (UniProt ID: P37202), and SpDis3L2 (UniProt ID: O14040); *Saccharomyces cerevisiae* – ScDis3 (UniProt ID: Q08162); *Caenorhabditis elegans* – CeDis3 (UniProt ID: Q17632), and CeDis3L2 (UniProt ID: Q09568); *Drosophila melanogaster* – DmDis3 (UniProt ID: Q9VC93), and DmDis3L2 (UniProt ID: Q8IRJ7); *Danio rerio* – DrDis3 (UniProt ID: E7FE01), DrDis3L1 (UniProt ID: A2RV18), and DrDis3L2 (UniProt ID: E7F350); *Xenopus tropicalis* – XtDis3 (UniProt ID: A0A6I8R570), XtDis3L1 (UniProt ID: Q0P4R5), and XtDis3L2 (UniProt ID: A0A6I8R0A9); *Gallus gallus* – GgDis3 (UniProt ID: A0A8V0X405), GgDis3L1 (UniProt ID: A0A8V0Z3V0), and GgDis3L2 (UniProt ID: A0A8V0Z2M1); *Mus musculus* – MmDis3 (UniProt ID: Q9CSH3), MmDis3L1 (also named mDis3L1, UniProt ID: Q8C0S1-1), and MmDis3L2 (UniProt ID: Q8CI75-1); *Homo sapiens –* HsDis3 (UniProt ID: Q9Y2L1-1), HsDis3L1 (UniProt ID: Q8TF46-1), and HsDis3L2 (UniProt ID: Q8IYB7-1); *Arabidopsis thaliana* – AtDis3 (UniProt ID: Q9SHL7), and AtDis3L2 (also named SOV, UniProt ID: P0DM58). The final figure was built using ESPript 3.0 v.4.0.26 (Robert and Gouet 2014), available at https://espript.ibcp.fr/ESPript/cgi-bin/ESPript.cgi.

### Cloning and expression of SpDis3L2 variants

The gene encoding SpDis3L2, located in plasmid pGEX-4T-1_SpDis3L2_WT (Malecki et al. 2013) was subcloned into a pET15b vector with an N-terminal His-SUMO tag. The point mutations D461N, C560Y, and A756R were introduced into pET15b_SpDis3L2_WT by overlapping PCR (primers listed in **Table S1**). All constructs were verified by DNA sequencing (StabVida, Portugal).

All plasmids (**Table S2**) were transformed into *E. coli* BL21 (DE3) CodonPlus – RIL cells (Stratagene) to produce the recombinant proteins. *E. coli* cultures were grown in TB medium (GRiSP, Boca Scientific) supplemented with 100 μg/ml ampicillin and 50 μg/ml chloramphenicol at 30°C. When cultures reached an OD_600_ of 0.5-0.8, protein expression was induced by adding 0.5 mM IPTG and cells were incubated at 16°C overnight. Cells were harvested by centrifugation and stored at –20°C.

### Purification of SpDis3L2 variants

Cells were resuspended in a buffer containing 100 mM MOPS pH 7.6, 200 mM KCl, and 1 mM PMSF, and lysed using the French Press (Thermo Electron). 5 U/ml benzonase nuclease (Sigma-Aldrich, Merck) were added, and lysates were clarified by centrifugation at 18,000 rpm at 4°C for 1 h.

Using the cleared lysate, a Ni-NTA affinity chromatography was performed in a HisTrap HP column (Cytiva). Proteins were eluted in a buffer containing 100 mM MOPS pH 7.6, 200 mM KCl, and 500 mM imidazole. Fractions were analyzed by 4-12% Tris-Gly SDS-PAGE, followed by BlueSafe (NZYTech) staining. After the protein fractions were pooled and concentrated, size exclusion chromatography was performed on a Superdex 200 Increase 10/300 GL column (Cytiva) pre-equilibrated in a buffer containing 100 mM MOPS pH 7.6, and 200 mM KCl. The collected protein fractions were analyzed by 4-12% Tris-Gly SDS-PAGE, followed by BlueSafe (NZYTech) staining. The fractions with the highest purity were pooled and concentrated. 50% (v/v) glycerol was added to the final protein samples prior to storage at –20 °C. All SpDis3L2 protein versions were purified at least twice in independent experiments to ensure reproducibility of results.

Purified proteins were quantified using the Bradford method (Bradford 1976), and full-length SpDis3L2 protein concentration in each sample was normalized by Western blot, using anti-6xHis tag (Invitrogen) as the primary antibody, and anti-rat IgG (Invitrogen) as the secondary antibody.

### Preparation of RNA substrates

30-mer poly(U), 20-mer Adh1, 24-mer Adh1-U_4_, and 36-mer Adh1-U_16_ (**Table S3**) synthetic oligoribonucleotides were used as substrates in the exoribonucleolytic activity and electrophoretic mobility shift assays. Each oligoribonucleotide was labelled at the 5’-end with [^32^P]-γ-ATP and T4 Polynucleotide Kinase (PNK, Ambion) for 1 h at 37°C. PNK was inactivated for 5 min at 80°C, and RNA oligomers were purified using illustra MicroSpin G-25 columns (Cytiva) to remove the non-incorporated nucleotides. To prepare the double-stranded RNA substrates, the radioactively labelled Adh1-U_16_ was hybridised to the complementary non-labelled 20-mer asAdh1 (**Table S3**) to obtain the corresponding Adh1-U_16_:asAdh1 dsRNA (dsAdh1-U_16_). The hybridisation was performed in a 1:5 molar ratio in 20 mM Tris-HCl pH 7.5 for 5 min at 80°C, followed by 60 min at RT.

To confirm the complete hybridisation of the molecules, the hybridisation mixture was resolved in a 20% (v/v) polyacrylamide (PAA) non-denaturing gel, in parallel with the respective ss form (Adh1-U_16_). Migration was compared through visualisation of the radioactive signal by PhosphorImaging (FujiFilm FLA-5100 Fluorescent Image Analyser).

### Exoribonucleolytic activity assays

Since SpDis3L2 is a 3’-5’ exoribonuclease, we used RNA substrates labelled at their 5’-ends in the exoribonucleolytic assays to monitor the formation of reaction products. The assays were carried out in a final reaction volume of 50 μl containing 8 nM labelled RNA substrate(s) and activity buffer (10 mM Tris-HCl pH 7.5, 25 mM NaCl, 5 mM MgCl_2_, and 1 mM DTT). Reactions were started by the addition of 20 nM of protein followed by incubation at 30°C. The protein variants were always tested in parallel with the WT protein, at the same concentration and from the same purification round. Control reactions, without the enzyme, were incubated under the same conditions.

Aliquots of 10 μl were withdrawn at different time points (indicated in the respective figure) and reactions were stopped by the addition of 5 μl gel-loading buffer containing 95% (v/v) formamide, 10 mM EDTA, 0.025% (w/v) bromophenol blue, and 0.025% (w/v) xylene cyanol. Reaction products were resolved in a 20% (v/v) polyacrylamide/7 M urea denaturing gel, and the radioactive signal was visualized by PhosphorImaging (FujiFilm FLA-5100 Fluorescent Image Analyser). Consumption of substrate over time was quantified using ImageQuant software (Cytiva). Each experiment was performed at least three times using proteins from two independent purifications.

### Electrophoretic mobility shift assay (EMSA)

Electrophoretic mobility shift assays (EMSAs) were carried out in a final volume of 10 μl containing 10 nM Adh1-U_16_, binding buffer (10 mM Tris-HCl pH 7.5, 25 mM NaCl, 1 mM DTT, and 50 mM EDTA), and increasing concentrations of protein (from 50 to 1000 nM). Reactions were incubated for 10 min at 30°C. Control reactions, without the enzyme, were incubated under the same conditions.

Then, 3 μl of gel-loading buffer containing 30% (v/v) glycerol, 0.25% (w/v) bromophenol blue, and 0.25% (w/v) xylene cyanol were added to the reaction. The RNA-protein complexes were subjected to ultraviolet crosslinking in a UVC 500 Crosslinker (UV Crosslinker, Amersham Biosciences, Cytiva). Samples were resolved in a 5% (v/v) polyacrylamide non-denaturing gel and the radioactive signal was visualised by PhosphorImaging (FujiFilm FLA-5100 Fluorescent Image Analyser).

## Data availability

The data underlying this article are available in the article and its online supplementary material. The atomic coordinates and structure factors for SpDis3L2–U_13_ have been deposited in the Protein Data Bank under accession codes 9CY7.

## Acknowledgements

We thank the support of the beamline staff at National Synchrotron Light Source II (FMX-17-ID-2), a U.S. Department of Energy (DOE) Office of Science User Facility operated for the DOE Office of Science by Brookhaven National Laboratory under Contract No. DE-SC0012704. We thank Teresa B. Silva for her technical support at ITQB NOVA.

## Author Contributions

RGM, CMA and SCV designed the study. RGM, AG, SMC, PP and SCV performed the experiments. AG and LJ performed the structural studies (experiments and analysis). RGM, AG, SMC and SCV analyzed the results and wrote the manuscript. SMC prepared the final version of the manuscript. All authors revised and approved the final version of the manuscript.

