## Supplemental Materials for "Structural and mechanistic insights into Dis3L2–mediated degradation of structured RNA"

### Supplemental Material

**Table S1.** Primers used in this study.

| Primer Name | Sequence 5' → 3' | Observations |
|---|---|---|
| T7Fw | TAATACGACTCACTATAGG | |
| T7Rev | GCTAGTTATTGCTCAGCGG | |
| 168_Fw | GCAGGCGAGAATCTTTATTTTCAGGGATCCTCATC CAATGCGAAAAAGTCAG | Primers for SLIC cloning SpDis3L2$^{\Delta N168}$ in pQlinkH. |
| 927_Rev | AAGAAAGCTGGGTCCTAGGCGGCCGCATTATTCA AAGAAACTAGACTACAGCG | |
| C560Y_Fw | CAGTAATAAAGACATATGCC | Primers for the introduction of C560Y mutation in the SpDis3L2 gene. |
| C560Y_Rev | GGCATATGTCTTTATTACTG | |
| A756R_Fw | GTGAGAAGACAGACTGGCACCATTACCGTCTTTC | Primers for the introduction of A756R mutation in the SpDis3L2 gene. |
| A756R_Rev | GAAAGACGGTAATGGTGCCAGTCTGTCTTCTCAC | |
| R458T_Fw | CCTGAAACAGCTACTGACTTGGATG | Primers for the introduction of R458T mutation in the SpDis3L2 gene. |
| R458T_Rev | CATCCAAGTCAGTAGCTGTTTCAGG | |

The bases that were changed are underlined.

**Table S2.** Plasmids used in this study.

| Plasmid Name | Source | Marker | Observations |
|---|---|---|---|
| pGEX-4T-1 _SpDis3L2_WT | (Malecki et al., 2013) | Amp$^R$ | Encodes wild-type full-length SpDis3L2 with a GST tag. |
| pGEX-4T-1 _SpDis3L2_D461N | (Malecki et al., 2013) | Amp$^R$ | Encodes a mutant full-length SpDis3L2 with a GST tag, where aspartic acid (D) at position 461 was substituted by an asparagine (N) (catalytically dead mutant). |
| pQlinkH_SpDis3L2 _ΔN168_D461N | This study | Amp$^R$ | Encodes a mutant SpDis3L2, where aspartic acid (D) at position 461 was substituted by an asparagine (N) (catalytically dead mutant) (Malecki et al., 2013). The gene has a ΔN168 N-terminal truncation, a N-terminal TEV cleavable 7xHis tag, and a C-terminal extra Asn-Ala-Ala-Ala tag. |
| pET15b_SpDis3L2 _WT | This study | Amp$^R$ | Adapted from pGEX-4T-1_GST_SpDis3L2_WT (Malecki et al., 2013); encodes wild-type full-length SpDis3L2 with a N-terminal His-SUMO tag (6xHis-SpDis3L2). |
| pET15b_SpDis3L2 _C560Y | This study | Amp$^R$ | Encodes a mutant 6xHis-SpDis3L2, where cysteine (C) at position 560 was substituted by a tyrosine (Y). |
| pET15b_SpDis3L2 _A756R | This study | Amp$^R$ | Encodes a mutant 6xHis-SpDis3L2, where alanine (A) at position 756 was substituted by an arginine (R). |

**Amp$^R$**: marker of resistance to ampicillin (*bla* gene encoded in the plasmid).

**Table S3.** RNA substrates used in this study for the crystallization, and for the exoribonucleolytic and electrophoretic mobility shift assays. All RNA substrates are synthetic oligoribonucleotides.

| RNA Substrate Name | Sequence 5' → 3' | Observations |
|---|---|---|
| $U_{13}$ | UUU UUU UUU UUU U | 13-nt sequence |
| Poly(U) | UUU UUU UUU UUU UUU UUU UUU UUU UUU UUU | 30-nt sequence |
| Adh1 | GUU UUG UAU AGA AAU CAA UG | 20-nt sequence at the 3'-end of *adh1* (alcohol dehydrogenase) mRNA from *S. pombe*; corresponds to Substrate 1 from (Malecki et al., 2013) |
| Adh1-$U_4$ | GUU UUG UAU AGA AAU CAA UGU UUU | 24-nt sequence; Adh1 sequence followed by 4 additional uridines |
| Adh1-$U_{16}$ | GUU UUG UAU AGA AAU CAA UGU UUU UUU UUU UUU UUU | 36-nt sequence; Adh1 sequence followed by 16 additional uridines |
| asAdh1 | CAU UGA UUU CUA UAC AAA AC | 20-nt antisense sequence complementary to Adh1 |

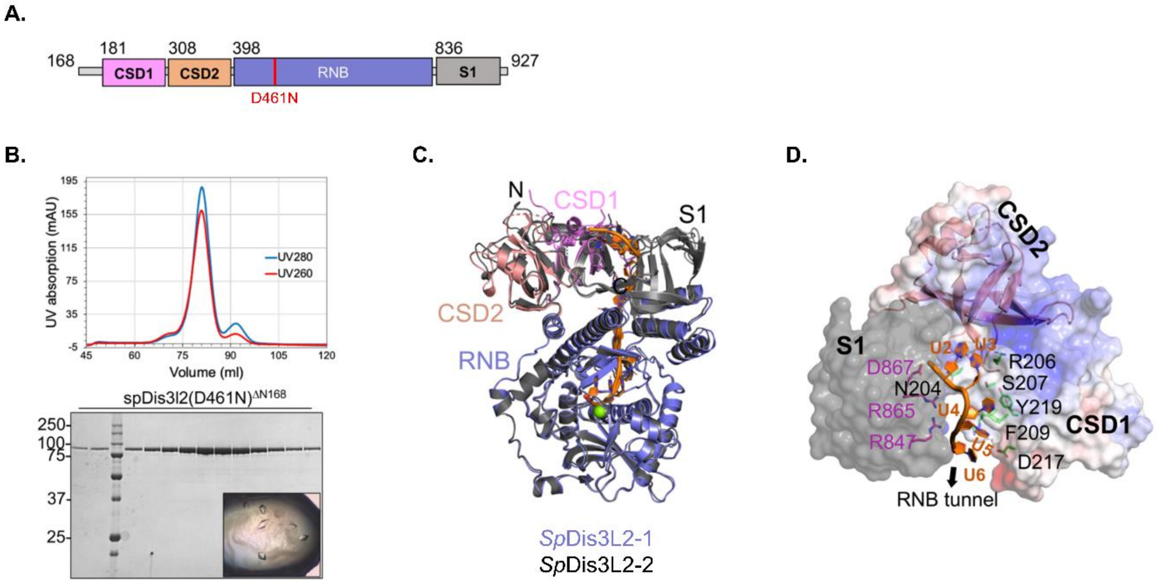


**Figure S1. Purification and structure determination of SpDis3L2. (A)** Domain architecture of SpDis3L2, highlighting CSD1, CSD2, RNB and S1 domains. **(B)** Size exclusion chromatogram and SDS-PAGE for purified SpDis3L2(D461N)$^{\Delta N168}$ protein used for crystallisation. The crystals used for structure determination are shown. **(C)** The superposition of the two molecules in the asymmetric unit of SpDis3L2(D461N)$^{\Delta N168}$–$U_{13}$ crystal structure. **(D)** The RNA interacts with CSD1, CSD2 and the S1 domain. The CSD surface's electrostatic potential is shown, while the S1 domain is shown in grey. Different residues interacting with nucleotides are also shown with sticks.

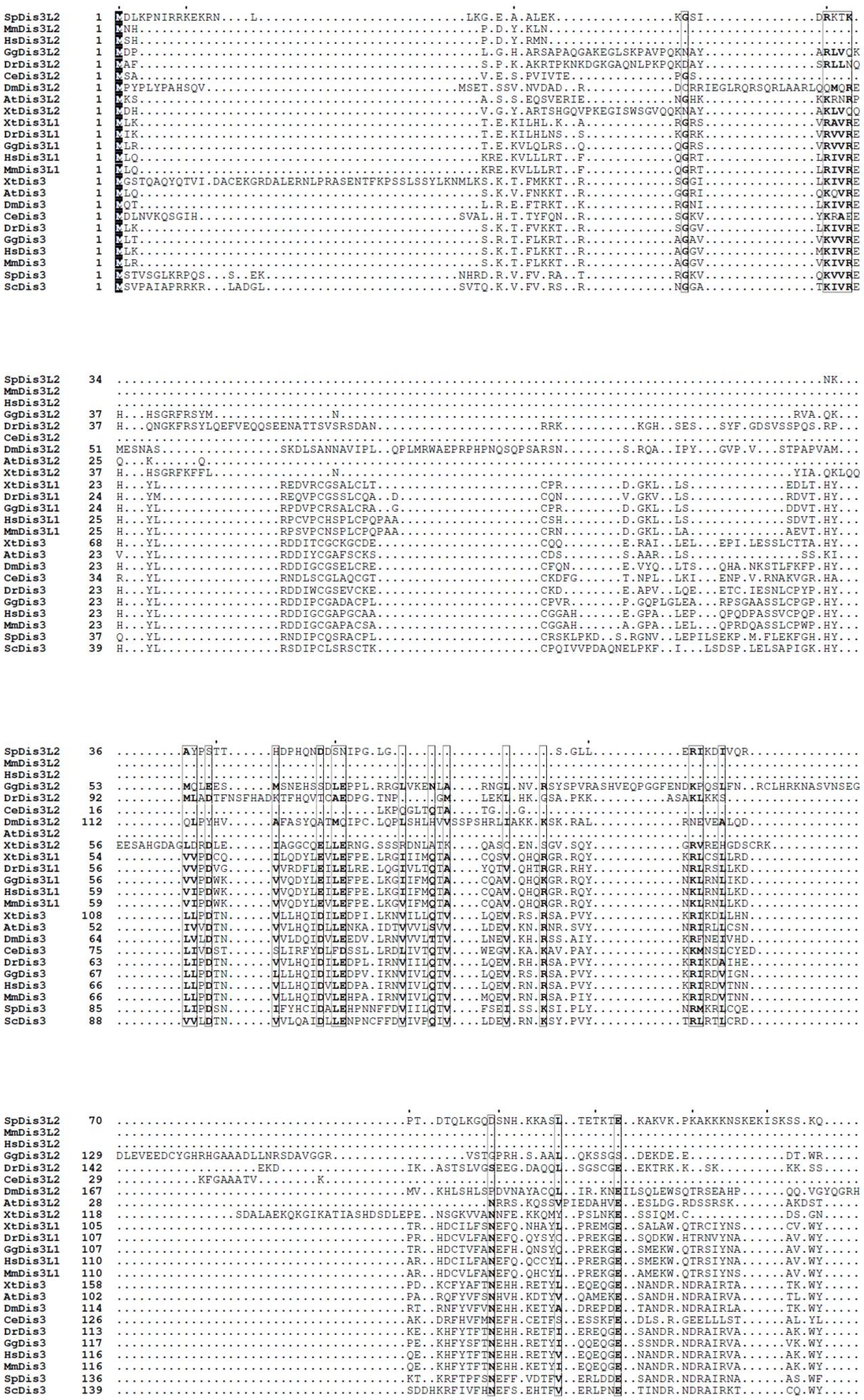

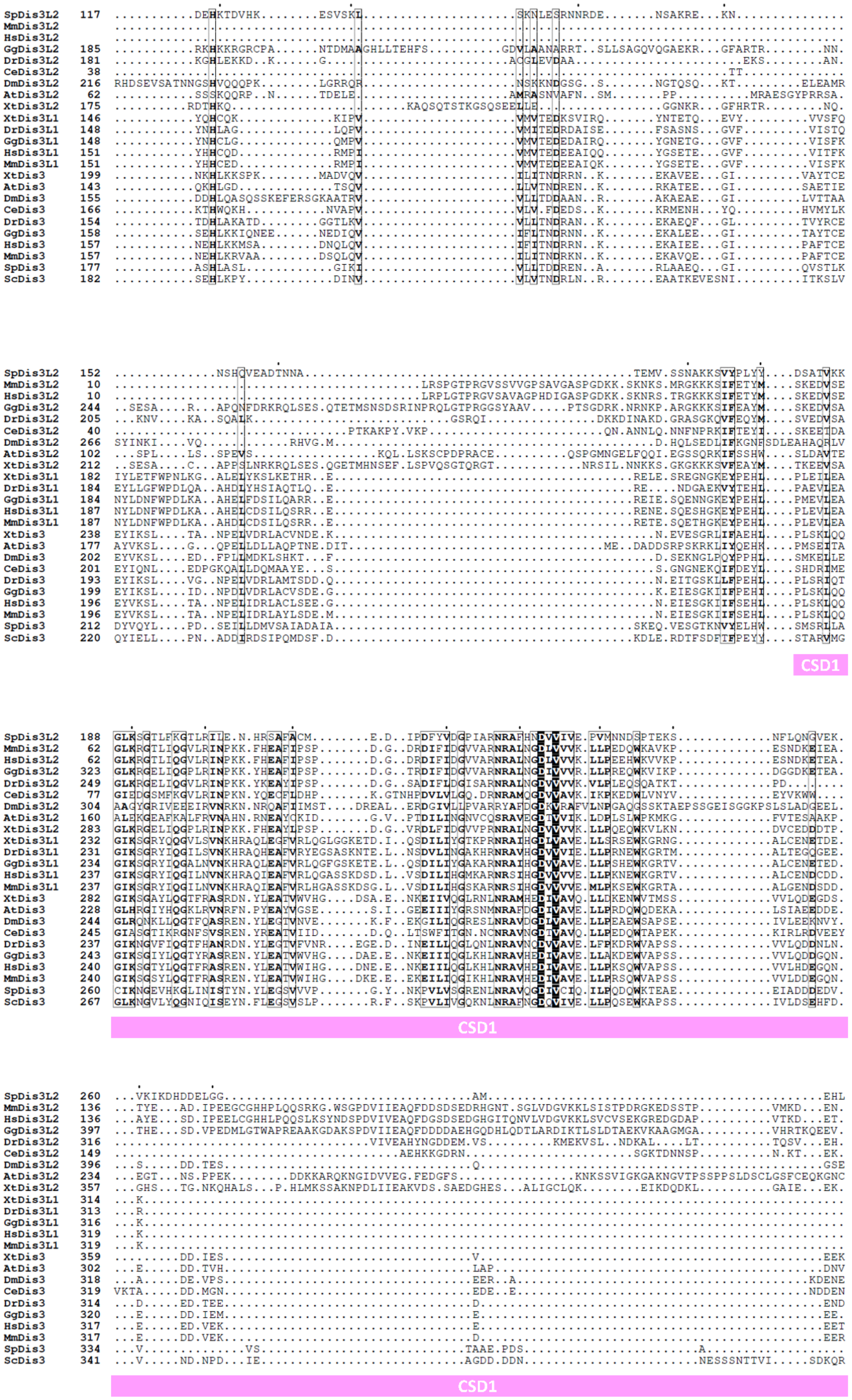
CSD1
CSD1
CSD1

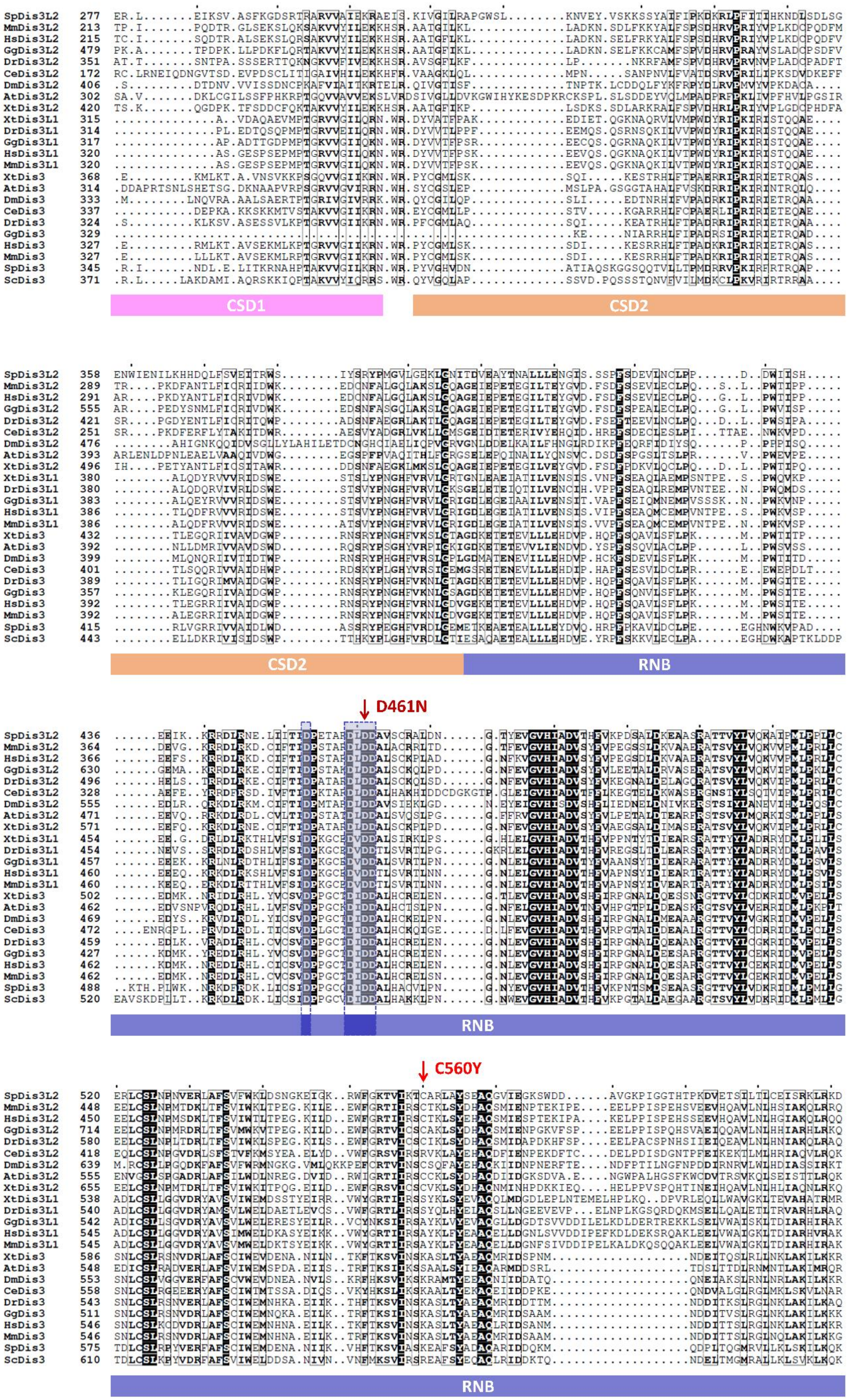
CSD1
CSD2
CSD2
RNB
D461N
RNB
C560Y
RNB

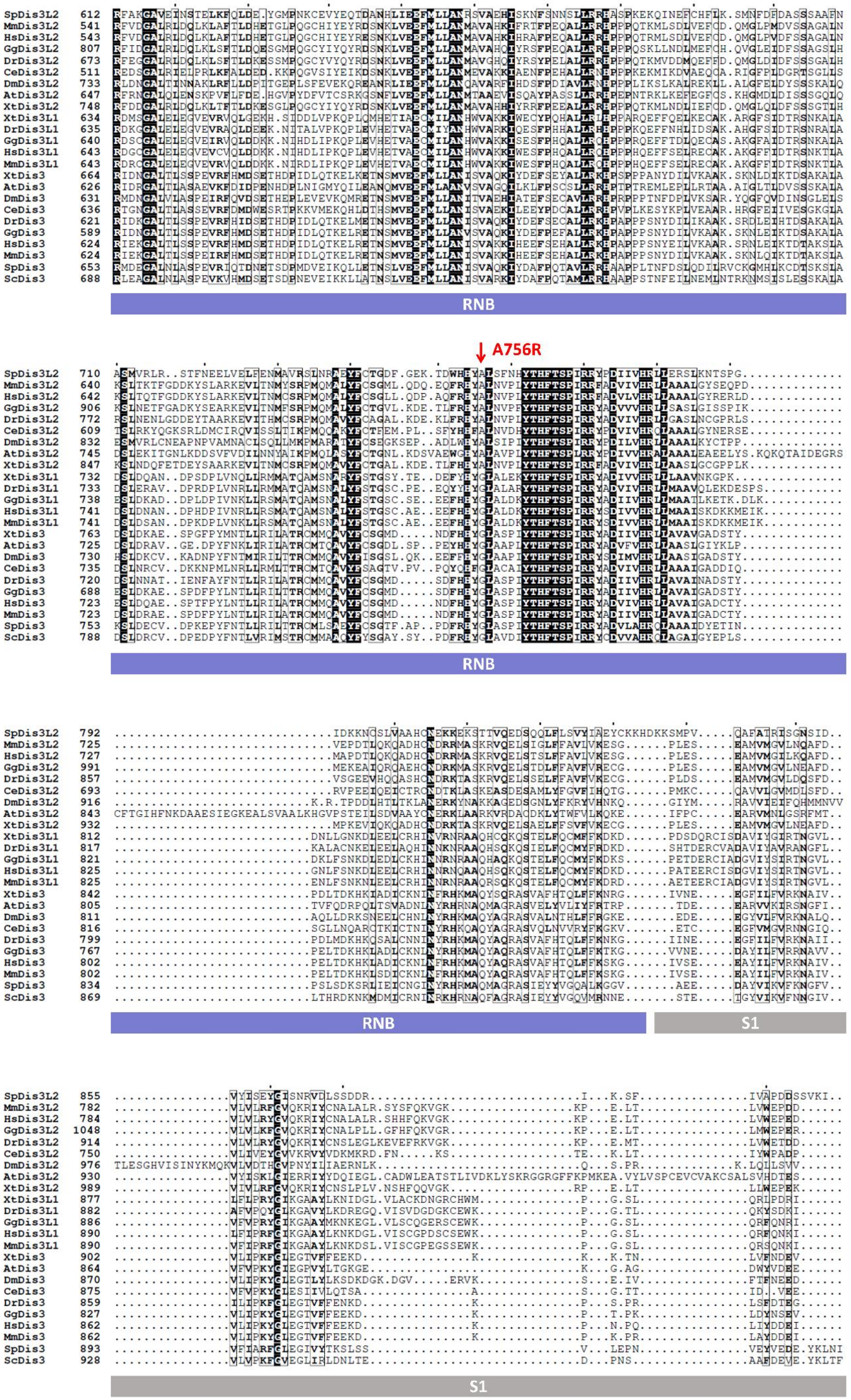
A756R
RNB
RNB
RNB
S1
S1

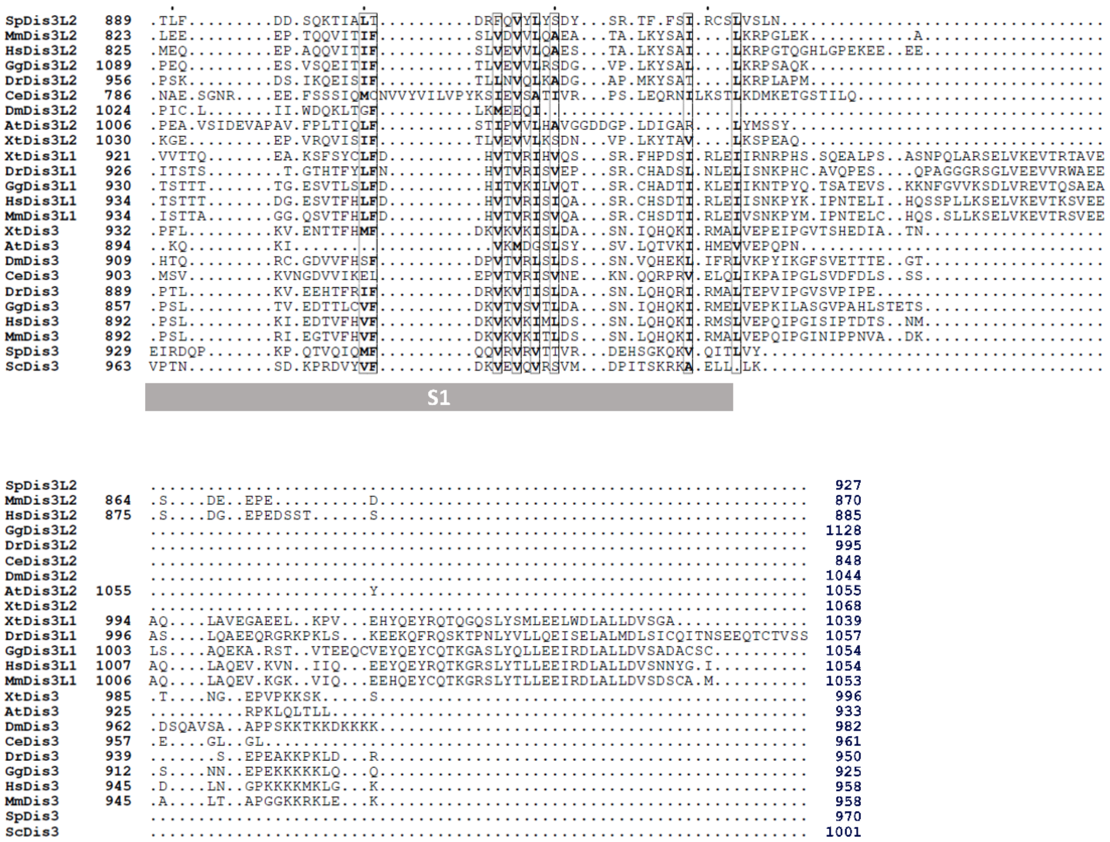


**Figure S2. Full-length sequence alignment of several eukaryotic homologues of the RNase II/RNB family of exoribonucleases, highlighting mutated residues.** The multiple sequence alignment was built using the default T-Coffee parameters and the amino acid sequences of the protein homologues of the RNase II/RNB family from the following species: *Schizosaccharomyces pombe* (fission yeast) – SpDis3 (also named Rrp44, UniProt ID: P37202), and SpDis3L2 (UniProt ID: O14040); *Saccharomyces cerevisiae* (budding yeast) – ScDis3 (also named Rrp44, UniProt ID: Q08162); *Caenorhabditis elegans* (worm) – CeDis3 (also named Rrp44, UniProt ID: Q17632), and CeDis3L2 (UniProt ID: Q09568); *Drosophila melanogaster* (fruit fly) – DmDis3 (UniProt ID: Q9VC93), and DmDis3L2 (UniProt ID: Q8IRJ7); *Danio rerio* (zebrafish) – DrDis3 (UniProt ID: E7FE01), DrDis3L1 (UniProt ID: A2RV18), and DrDis3L2 (UniProt ID: E7F350); *Xenopus tropicalis* (frog) – XtDis3 (UniProt ID: A0A6I8R570), XtDis3L1 (UniProt ID: Q0P4R5), and XtDis3L2 (UniProt ID: A0A6I8R0A9); *Gallus gallus* (chicken) – GgDis3 (UniProt ID: A0A8V0X405), GgDis3L1 (UniProt ID: A0A8V0Z3V0), and GgDis3L2 (UniProt ID: A0A8V0Z2M1); *Mus musculus* (mouse) – MmDis3 (also named mDis3 or Rrp44, UniProt ID: Q9CSH3), MmDis3L1 (also named mDis3L1, UniProt ID: Q8C0S1-1), and MmDis3L2 (also named mDis3L2, UniProt ID: Q8CI75-1); *Homo sapiens* (human) – HsDis3 (also named hDIS3 or RRP44, UniProt ID: Q9Y2L1-1), HsDis3L1 (also named hDIS3L1, UniProt ID: Q8TF46-1), and HsDis3L2 (also named hDIS3L2, UniProt ID: Q8IYB7-1); *Arabidopsis thaliana* (flowering plant) – AtDis3 (also named RRP44A, UniProt ID: Q9SHL7), and AtDis3L2 (also named SOV, UniProt ID: P0DM58). This final figure was constructed using ESPript 3.0 v.4.0.26. Residues are colour-coded in grey scale according to their amino acid conservation: black boxes are positions with identical residues in all proteins; white boxes with bold lettering are positions with residues with similar properties. The coloured rectangular boxes under the alignment indicate in which protein domain each region is located, using SpDis3L2 positions and colour-coded as in **Figure 1.** The two dark blue boxes in the RNB domain denote the four aspartic acid residues (D) of the active site of the enzymes, which are conserved in the RNB domain and essential for its exonucleolytic function. The second dark blue box includes the key residue that distinguishes the amino acid signature of the catalytic center of the three paralogues: DLDD for Dis3L2, DIDD for Dis3, and DVDD for Dis3L1 proteins. A comprehensive analysis of different consensus sequences for the catalytic residues of Dis3-like proteins was recently published, showing that there are exceptions, namely, many Dis3L1 proteins have a DIDD catalytic motif (Ballou *et al.*, 2021; *Mol. Biol. Evol.* 38(5):1837-1846; doi: 10.1093/molbev/msaa324). The positions modified in SpDis3L2 in this work are outlined in red, and the respective substitutions are referred above the respective residues – D461N (catalytically inactive), C560Y, and A756R SpDis3L2 variants. The C560Y SpDis3L2 mutant mimics the C489Y hDIS3L2 mutant found in a patient with Perlman Syndrome. The A756R SpDis3L2 mutant mimics the G766R hDIS3 mutant found in a patient with Multiple Myeloma.

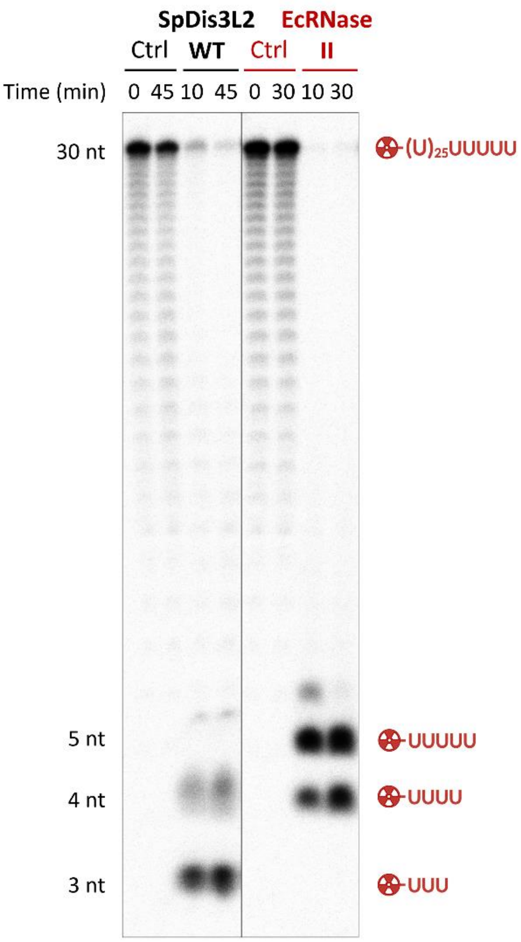


**Figure S3. Exoribonucleolytic activity of *S. pombe* Dis3L2 (SpDis3L2) and *E. coli* RNase II (EcRNase II) over a poly(U) RNA substrate – comparison of end-product size.** 8 nM of poly(U) substrate was incubated either with 30 nM WT SpDis3L2 for 45 minutes at 30ºC, or with 40 nM EcRNase II for 30 minutes at 37ºC. Control reactions (Ctrl) were incubated for the same time without any enzyme. Reactions were stopped at the time points indicated above each lane. RNA substrates and degradation products were separated through migration in a 7 M urea/20% PAA denaturing gel. The sizes of the molecules are depicted alongside the gel in line with the corresponding bands.

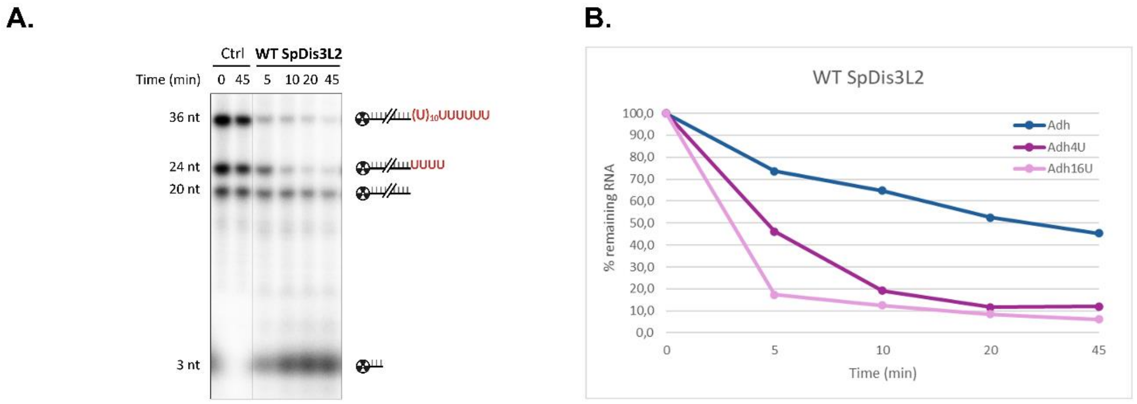


**Figure S4. Exoribonucleolytic activity of wild-type SpDis3L2 on three ssRNA, demonstrating its preference for uridylated RNA substrates.** Adh1 (20 nt), Adh1-$U_4$ (24 nt) and Adh1-$U_{16}$ (36 nt) RNA substrates were simultaneously incubated with wild-type (WT) SpDis3L2 for 45 minutes at 30ºC. Control reactions (Ctrl) were incubated for the same time without any enzyme. Reactions were stopped at the time points indicated above each lane and analysed on a **7 M urea/20% PAA denaturing gel (A)**. The sizes of the molecules are depicted alongside the gel in line with the corresponding bands. This assay was performed more than three times. **(B) Quantification of the exoribonucleolytic activity of WT SpDis3L2.** The graph represents the amount of remaining RNA at the time points indicated, for each of the three RNA substrates (Adh1, Adh1-$U_4$, and Adh1-$U_{16}$). The amount of remaining RNA was quantified using the ImageQuant TL v.8.1 software.
